# Kinetochore Proteins CENP-B and LIC2 are critical for Episome Persistence and Replication of Kaposi’s Sarcoma-Associated Herpesvirus

**DOI:** 10.64898/2026.10.28.762626

**Authors:** Rajnish Kumar Singh, Shengwei Zhang, Dipayan Bose, Nian Ma, Ramakrishna Vangala, Kiran Chunduru, Erle S. Robertson

## Abstract

Kaposi’s sarcoma-associated herpesvirus (KSHV) establishes lifelong latency by maintaining its genome as circular episomes tethered to host chromatin and faithfully segregated during mitosis. This process is primarily mediated by the latency-associated nuclear antigen (LANA), which interacts with host chromosome-associated factors to ensure viral genome persistence. Several kinetochore proteins, including CENP-F, Bub1, BubR1, Aurora B, Survivin, and NuMA, have been implicated in this process, yet the broader contribution of kinetochore components remains poorly defined. We performed a systematic shRNA-based screen of the 37 host genes encoding kinetochore-associated proteins to evaluate their role in KSHV episome maintenance. We validated involvement of previously reported factors and identified Centromere Protein B (CENP-B) and Light Intermediate Chain 2 of the dynein complex (LIC2) as novel kinetochore-associated proteins critical for KSHV episome persistence. Loss of these proteins significantly reduced episome stability, highlighting previously unrecognized host-virus interactions at the chromosome segregation machinery. These findings identify critical host factors that expand the repertoire of kinetochore proteins co-opted by KSHV for episome maintenance and provide new mechanistic insights into how viral episomes exploit host chromosome segregation pathways for long-term persistence. Our work underscores kinetochore proteins as a broader class of host dependency factors in Ƴ-herpesvirus biology and suggests potential therapeutic targets for KSHV-associated malignancies.

## Introduction

Kaposi’s sarcoma-associated herpesvirus (KSHV; human herpesvirus 8) is an oncogenic γ-herpesvirus etiologically associated with Kaposi’s sarcoma, primary effusion lymphoma (PEL), and KSHV-associated multicentric Castleman disease[1–4]. A defining feature of KSHV infection is its ability to establish long-term latency, during which the viral genome persists predominantly as a circular, extrachromosomal episome within the nucleus of infected cells[5–8]. Unlike integrated viral genomes, latent KSHV episomes must independently overcome two fundamental challenges during proliferation of the host cell: the viral DNA must be efficiently duplicated and the replicated episomes must be faithfully distributed to daughter nuclei during cell division. Failure of either process progressively reduces viral genome copy number and ultimately compromises persistence of the latent reservoir[9, 10].

The latency-associated nuclear antigen (LANA) is central to both processes of KSHV replication and segregation[11, 12]. LANA binds sequence-specific sites within the KSHV terminal repeats (TRs) through its C-terminal DNA-binding domain, while regions within its N terminus associate with host chromatin, thereby tethering viral episomes to cellular chromosomes[13–16]. This chromosome-tethering function enables KSHV genomes to remain associated with host chromatin throughout mitosis and facilitates their inheritance by daughter cells[17, 18]. LANA and the TR region also provide an important platform for latent viral DNA replication through recruitment or engagement of cellular replication factors[19–21]. Accordingly, disruption of LANA expression, LANA-TR binding, or chromosome tethering results in progressive loss of KSHV episomes[22, 23]. LANA therefore provides a molecular interface through which an extrachromosomal viral genome exploits host chromosome biology for its long-term maintenance.

Although LANA is the principal viral determinant of episome persistence, faithful inheritance of KSHV genomes cannot be accomplished by a viral tether alone. Host chromosomes undergo highly coordinated structural and spatial changes as cells enter mitosis, align their chromosomes at the metaphase plate, segregate sister chromatids, and re-establish daughter nuclei[24, 25]. KSHV episomes associated with these chromosomes must remain compatible with this dynamic process[15, 16]. Increasing evidence suggests that LANA and KSHV episomes engage cellular proteins involved in centromere organization, kinetochore function, spindle attachment, and mitotic regulation[16, 26, 27]. Individual factors including CENP-F, Bub1, BubR1, Aurora B, Survivin, NuMA, and other chromosome-associated proteins have been implicated in LANA localization or viral episome maintenance [18, 28–32]. These observations raise the possibility that KSHV persistence depends not simply on attachment to chromatin, but on coordinated engagement of the cellular machinery that regulate chromosome behavior during cell division.

The kinetochore is a large multiprotein assembly formed at centromeric chromatin that provides the principal interface between chromosomes and spindle microtubules[33]. The Kinetochore architecture comprises an inner kinetochore assembled on centromeric chromatin and an outer kinetochore that establishes microtubule attachment, together with regulatory networks that monitor attachment, tension, and chromosome segregation [33–35]. The constitutive centromere-associated network (CCAN) connects specialized centromeric chromatin to outer kinetochore complexes, whereas the KNL1–Mis12–NDC80 (KMN) network provides a major microtubule-binding interface[36–39]. These structural components function together with spindle-assembly checkpoint proteins, chromosomal passenger proteins, microtubule regulators, dynein-associated machinery, and Ran-dependent factors to ensure accurate chromosome alignment and segregation[40–42]. Disruption of these systems can result in chromosome misalignment, lagging chromosomes, abnormal DNA content, and genome instability[43–45].

Despite the intimate relationship between latent KSHV episomes and host chromosomes, the extent to which this broader kinetochore and chromosome-segregation network contributes to viral genome persistence remains incompletely understood[16, 27]. Previous studies have largely focused on individual host proteins, making it difficult to determine whether the reported interactions represent isolated mechanisms or components of a broader host dependency network[16, 46]. Moreover, the relationship may extend beyond physical partitioning of viral genomes during mitosis[47]. Whether kinetochore-associated factors influence replication of KSHV episomes in addition to their segregation therefore represents an important unresolved question.

A second unresolved issue is whether the relationship between KSHV and the kinetochore-associated machinery is unidirectional. KSHV may simply utilize constitutively available chromosome-associated proteins, or infection may alter components of this machinery to generate a cellular environment favorable for viral persistence. Transcriptomic remodeling is a prominent feature of KSHV infection and latency[7, 48], nevertheless, systematic examination of kinetochore and chromosome segregation-associated gene expression in KSHV-positive cells has been limited. Determining whether infection selectively alters these pathways, and whether individual viral proteins contribute to such changes, could distinguish passive utilization of host machinery from broader virus-associated remodeling of chromosome-regulatory networks.

Here, we systematically investigated the contribution of kinetochore- and chromosome segregation-associated proteins to KSHV episome persistence. Using an shRNA-based functional screen encompassing 37 candidate proteins, we identified host factors whose depletion altered viral genome maintenance and prioritized Centromere Protein B (CENP-B) and the Dynein light intermediate chain protein 2 (LIC2) for mechanistic investigation. Their requirement was subsequently examined in naturally infected KSHV-positive PEL cells, together with the relationship between KSHV infection and transcriptional remodeling of kinetochore-associated genes. Using complementary biochemical and imaging approaches, including co-immunoprecipitation, bimolecular fluorescence complementation, FRET, and cell-cycle imaging, we examined the association of these proteins with LANA and their relationship to mitotic chromosome behavior. Finally, single-molecule analysis of replicated DNA and ChIP-based analysis of LANA and PCNA occupancy at viral terminal repeats were used to investigate whether kinetochore-associated factors also contribute to episome replication. Together, these studies identify a broader functional connection between host kinetochore-associated machinery and KSHV genome persistence and suggest that viral episome inheritance involves coordinated engagement of the cellular mechanisms governing chromosome segregation and DNA replication.

## Results

### Kinetochore and chromosome segregation proteins contribute to the persistence of KSHV episomes

To systematically investigate the contribution of host kinetochore and chromosome segregation machinery to KSHV episome maintenance, we curated a candidate set of proteins representing multiple functional layers of the chromosome segregation apparatus, including components of the inner and outer kinetochore, spindle checkpoint and segregation machinery, and microtubule-associated factors (Fig. 1A).

**Figure 1.**
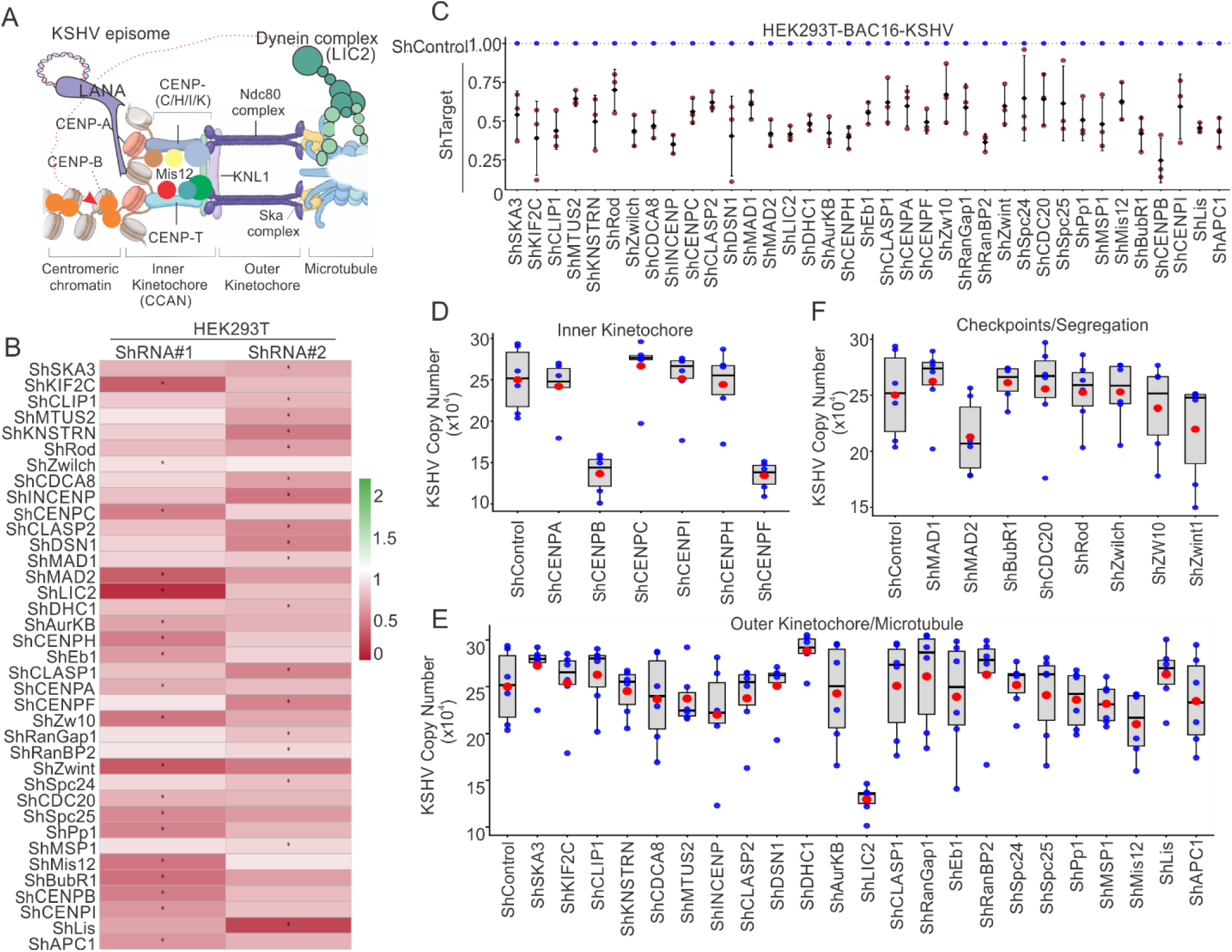
Systematic depletion of kinetochore and chromosome segregation proteins identifies host factors required for KSHV episome maintenance. (A) Schematic representation of kinetochore organization illustrating centromeric chromatin, the inner kinetochore/constitutive centromere-associated network (CCAN), and the outer kinetochore KMN network that connects centromeric chromatin to spindle microtubules. Representative proteins from the functional groups investigated in the shRNA screen are indicated. (B) Evaluation of shRNA-mediated depletion of kinetochore and chromosome segregation-associated genes. Two independent shRNA constructs targeting each candidate gene were evaluated in HEK293T cells, and transcript abundance was determined by quantitative real-time PCR. Knockdown efficiency is represented relative to the corresponding shControl condition. The more effective shRNA for each candidate was selected for subsequent functional analysis. (C) Knock-down confirmation of kinetochore and chromosome segregation-associated genes in HEK293T-BAC-KSHV cells. The more effective shRNA for each candidate was selected from HEK293T experiment was used to transfect HEK293T-BAC-KSHV cells. (D-F) Systematic analysis of KSHV genome copy number following depletion of the selected kinetochore and chromosome segregation-associated proteins in HEK293T-BAC16-KSHV cells. KSHV genome copy number was determined by quantitative PCR using equal amounts of genomic DNA and normalized to the common shControl reference and organized according to the principal functional classification of the targeted proteins. Candidates are grouped as inner kinetochore-associated proteins (D), outer kinetochore/microtubule-associated proteins (E), and checkpoint/chromosome segregation-associated proteins (F). Blue points represent individual measurements, and red points represent mean value. Box-and-whisker elements summarize the distribution of measurements for each condition. shControl is shown as the reference condition for comparison.

We first evaluated two independent shRNA constructs directed against each of the 37 candidate genes. Knockdown efficiency was initially assessed in HEK293T cells by quantitative real-time PCR. The two shRNA constructs produced varying degrees of transcript depletion across the candidate set, with at least one construct providing substantial knockdown for the majority of genes examined (Fig. 1B). Based on these measurements, the more effective shRNA for each target was selected for subsequent functional screening in HEK293T-BAC16-KSHV cells.

To determine whether depletion of kinetochore-associated genes affects the KSHV genome number, cells were transfected with the selected shRNAs (Fig. 1C) and viral genome copy number was quantified by absolute KSHV DNA copy-number analysis using equal amounts of genomic DNA. Comparison across the complete candidate set revealed a broad range of effects on viral genome abundance. Whereas depletion of several proteins produced relatively modest changes, knockdown of a subset of candidates resulted in pronounced reductions in KSHV copy number, indicating that maintenance of viral episomes is particularly sensitive to disruption of specific components of the chromosome segregation machinery (Fig. 1C-F).

To better understand whether these effects were associated with particular functional components of the kinetochore machinery, candidates were subsequently organized according to their principal roles in chromosome segregation. Among inner kinetochore-associated proteins, depletion of individual components produced markedly different effects on KSHV genome abundance, with several targets causing substantial reductions relative to the shControl condition (Fig. 1D). Similarly, depletion of proteins associated with the outer kinetochore and microtubule attachment machinery affected viral copy number to varying degrees (Fig. 1E). Perturbation of checkpoint and chromosome segregation-associated factors also altered KSHV genome abundance, although the magnitude of the phenotype differed considerably among individual proteins (Fig. 1F).

Together, this study demonstrates that KSHV episome maintenance is sensitive to perturbation of multiple components of the host kinetochore and chromosome segregation machinery. Importantly, the effects were not uniform across all candidates, suggesting that viral genome persistence does not simply depend on general kinetochore integrity but may exhibit a selective requirement for particular host kinetochore factors. Among the candidates producing prominent and reproducible effects, CENP-B and LIC2 emerged as proteins of particular interest and were therefore prioritized for subsequent mechanistic investigation.

### CENP-B and LIC2 contribute to persistence of KSHV episomes in naturally infected KSHV-positive pleural effusion lymphoma cells

To validate the functional relevance of candidate kinetochore-associated proteins identified in the HEK293T-BAC16-KSHV screen, we examined their contribution to viral episome maintenance in naturally infected KSHV-positive primary effusion lymphoma (PEL) cells. Based on the initial functional screen, CENP-B and LIC2 emerged as prominent candidates whose depletion substantially reduced KSHV genome abundance. We therefore selected these proteins for further validation in two naturally infected PEL-derived cell lines, BC3 and BCBL1.

Stable depletion of CENP-B or LIC2 was established by lentivirus mediated transduction of the corresponding shRNA constructs, with GFP serving as a marker of transduced cells. Fluorescence microscopy demonstrated robust GFP-positive populations in shControl, shCENPB, and shLIC2 transduced BC3 and BCBL1 cells (Fig. 2A,B). Quantitative real-time PCR confirmed significant depletion of the corresponding transcripts relative to shControl cells in both KSHV-positive cell backgrounds (Fig. 2C), establishing stable knockdown systems for subsequent analysis of viral episome maintenance.

**Figure 2.**
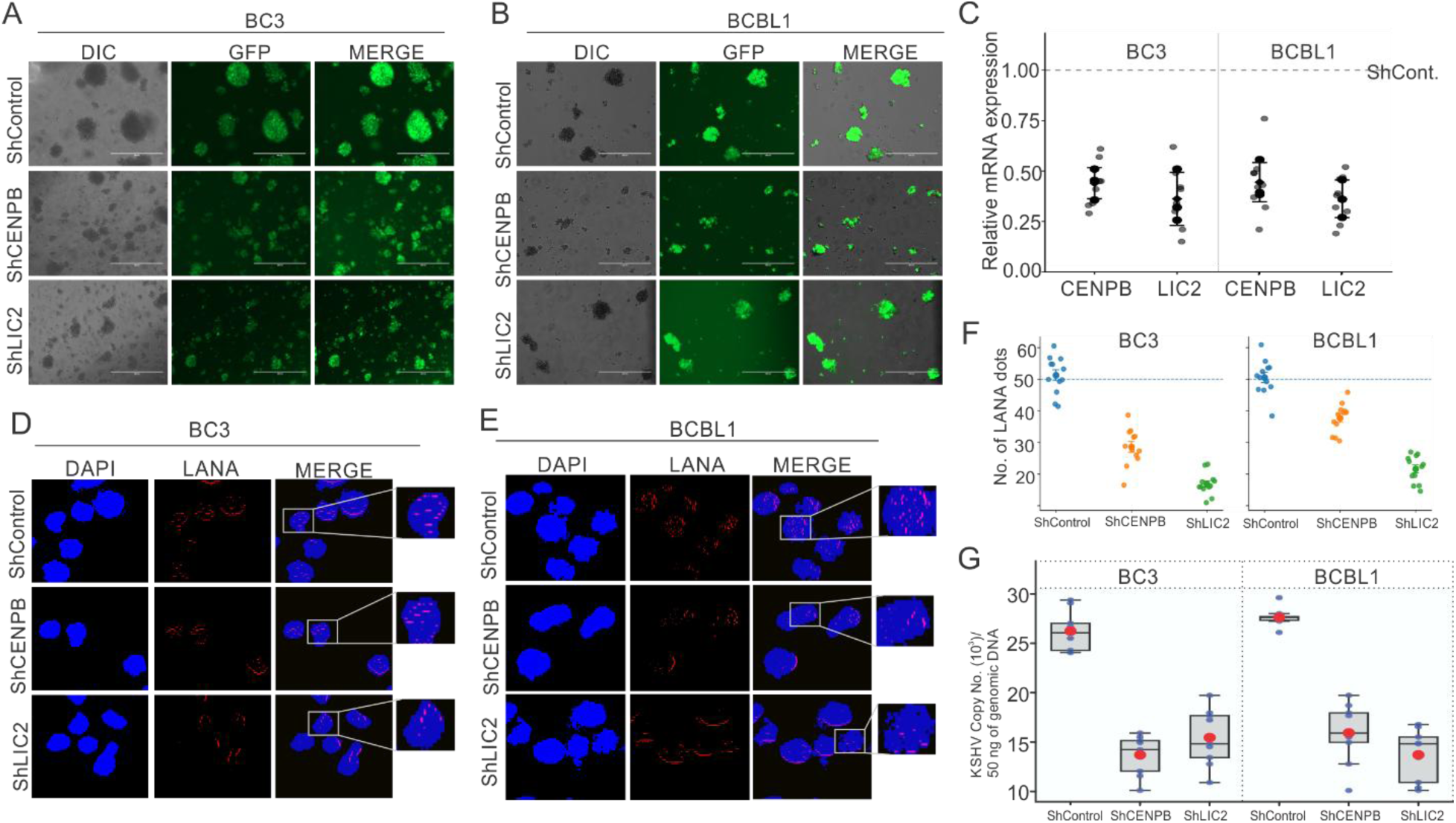
CENPB and LIC2 contribute to KSHV episome maintenance in naturally infected primary effusion lymphoma cells. (A, B) Establishment of stable CENPB and LIC2 depletion in naturally infected KSHV-positive PEL cell lines. Representative DIC, GFP fluorescence, and merged images of BC3 (A) and BCBL1 (B) cells stably transduced with shControl, shCENPB, or shLIC2 constructs. GFP expression indicates cells carrying the respective lentiviral shRNA constructs. (C) Quantitative real-time PCR analysis confirming depletion of CENPB and LIC2 transcripts in stable BC3 and BCBL1 knockdown cells. Transcript abundance was normalized to GAPDH and expressed relative to the corresponding shControl condition. (D, E) Representative immunofluorescence images showing LANA nuclear puncta in shControl, shCENPB, and shLIC2-expressing BC3 (D) and BCBL1 (E) cells. Cells were immune-stained for LANA (red), and nuclei were counterstained with DAPI (blue). Merged images show the distribution of LANA-associated nuclear foci under each condition. (F) Quantification of LANA nuclear puncta in individual BC3 and BCBL1 cells following depletion of CENPB or LIC2. Each point represents the number of LANA puncta detected in an individual cell. (G) Absolute quantification of KSHV genome copy number in BC3 and BCBL1 cells following depletion of CENPB or LIC2. Viral genome copy number was determined by quantitative PCR using KSHV-specific primers and normalized to cellular genomic DNA.

We next examined the effect of CENP-B and LIC2 knock-down on latent KSHV episomes at the single-cell level. Since LANA associates with KSHV terminal repeat sequences and forms characteristic punctate nuclear foci corresponding to chromosome-associated viral episomes[49], cells were examined by immunofluorescence microscopy for LANA nuclear puncta. Numerous discrete LANA foci were observed in nuclei of shControl BC3 and BCBL1 cells (Fig. 2D,E). In contrast, knock-down of either CENP-B or LIC2 resulted in a pronounced reduction in the number of LANA-associated nuclear puncta. Quantification across individual cells confirmed a substantial decrease in LANA foci following depletion of either protein in both cell backgrounds (Fig. 2F), consistent with reduced maintenance of latent KSHV episomes.

To independently quantify the effect on viral genome abundance, KSHV DNA copy number was measured by absolute quantitative PCR. Viral genome abundance was normalized to cellular genomic DNA to account for differences in cell number and DNA input. Consistent with the reduction in LANA nuclear puncta, knock-down of CENP-B or LIC2 resulted in decreased KSHV genome copy number in both BC3 and BCBL1 cells compared with their respective shControl cells (Fig. 2G). The concordant phenotypes observed across independent experiments and in two naturally infected PEL cell lines support a requirement for both CENP-B and LIC2 in efficient KSHV episome maintenance.

Collectively, these findings validate CENP-B and LIC as host factors contributing to KSHV episome persistence beyond the initial HEK293T-BAC16-KSHV screening system. The agreement between the functional screen, single-cell LANA-focus analysis, and absolute viral genome quantification demonstrates that depletion of either protein compromises maintenance of latent KSHV episomes in naturally infected PEL cells and provided the basis for investigating the mechanisms through which these kinetochore-associated factors support viral persistence.

### KSHV infection selectively modulates the expression of kinetochore-associated genes

Having established that multiple kinetochore-associated proteins contribute to KSHV episome maintenance, we next asked whether KSHV infection is accompanied by transcriptional remodeling of the host kinetochore machinery genes. To address this question, we analyzed a existing RNA-sequencing dataset generated from parental BJAB cells and BJAB cells stably maintaining KSHV episomes (BJAB-KSHV)[50]. From the transcriptome-wide dataset, we extracted a curated set of kinetochore and chromosome segregation-associated genes corresponding to the functional groups represented in our initial shRNA screen.

Comparison of BJAB-KSHV with parental BJAB cells revealed selective rather than uniform changes in the expression of kinetochore-associated genes (Fig. 3A). Differentially expressed genes were distributed across multiple functional components of the kinetochore and chromosome segregation machinery, including constitutive centromere/CCAN proteins, chromosomal passenger and mitotic regulatory factors, microtubule-associated proteins, and Ran-dependent chromosome segregation machinery (Fig. 3B). Examination of the overall distribution of expression changes further indicated that KSHV-associated transcriptional remodeling was restricted to a subset of kinetochore-associated genes rather than representing a coordinated global increase or decrease in this gene class (Fig. 3C). Notably, CENPB and LIC2, which emerged as prominent functional determinants of KSHV episome maintenance in our knockdown studies, were not significantly altered at the transcript level in the BJAB-KSHV RNA-seq dataset. Thus, the functional requirement for these proteins is separate from their transcriptional regulation by KSHV.

**Figure 3.**
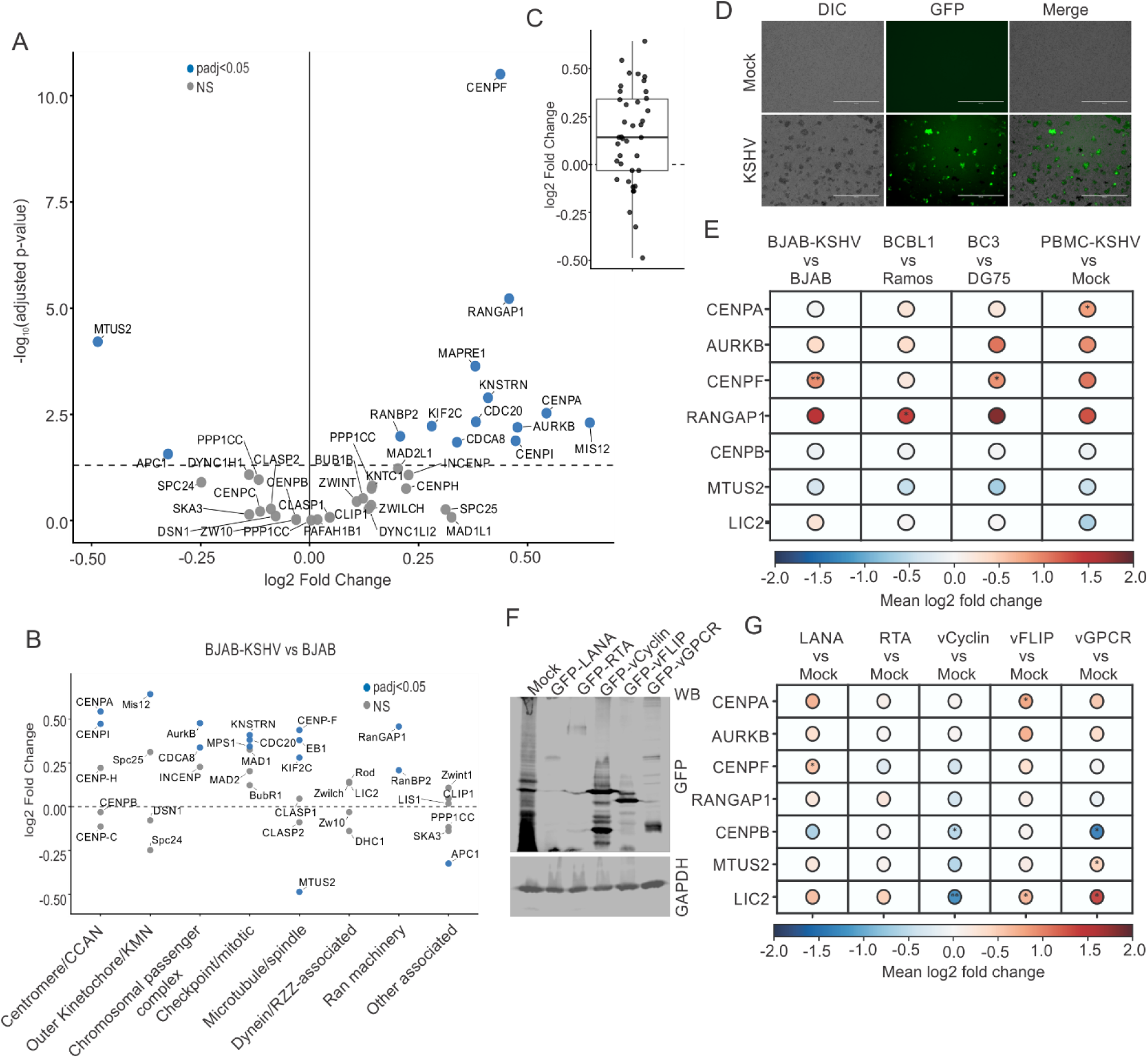
KSHV infection is associated with selective transcriptional remodeling of kinetochore-associated genes and identification of individual viral proteins. **(A)** Differential expression of kinetochore-associated genes in KSHV-positive BJAB-KSHV cells relative to parental BJAB cells determined by re-analysis of RNA-sequencing data. The scatter plot displays log2 fold change versus −log10 adjusted *P* value for the curated set of kinetochore and chromosome segregation-associated genes. Genes meeting the indicated significance criterion (adjusted *P* < 0.05) are highlighted. **(B)** Functional organization of kinetochore-associated transcriptional changes identified in BJAB-KSHV cells. Genes were grouped according to their major kinetochore and chromosome segregation-associated functions, including centromere/CCAN, outer kinetochore/KMN, chromosomal passenger complex, checkpoint/mitotic control, microtubule/spindle, dynein/RZZ-associated, Ran machinery, and other associated factors. Log2 fold changes are shown for individual genes, with significantly altered genes indicated in color. **(C)** Distribution of RNA-seq-derived log2 fold changes across the curated kinetochore-associated gene set, illustrating the overall magnitude and direction of transcriptional changes in BJAB-KSHV relative to BJAB cells. **(D)** Representative DIC, GFP fluorescence, and merged microscopy images of peripheral blood mononuclear cells (PBMCs) following mock or GFP-KSHV infection. GFP-positive cells indicate KSHV infection. **(E)** Cross-model quantitative real-time PCR analysis of selected kinetochore-associated genes. Transcript expression was examined in BJAB-KSHV relative to BJAB cells, BCBL1 relative to Ramos cells, BC3 relative to DG75 cells, and KSHV-infected relative to mock infected PBMCs. Selected genes include representative RNA-seq-responsive genes together with CENPB and LIC2, which were retained as functionally important comparators. Fold-change magnitude and statistical significance are represented according to the color and asterisks, respectively. **(F)** Immunoblot analysis confirming ectopic expression of individual KSHV proteins in HEK293T cells transfected with constructs encoding LANA, RTA, vCyclin, vFLIP, or vGPCR. GFP immunoblotting was used to detect the respective GFP-tagged viral proteins, and GAPDH served as a loading control. **(G)** Quantitative real-time PCR analysis of selected kinetochore-associated transcripts following ectopic expression of individual KSHV proteins in HEK293T cells. Changes in fold change expressions relative to mock-transfected cells are represented by color, while statistical significance is represented by asterisks.

To determine whether selected transcriptional changes identified by RNA-seq were also evident in additional KSHV-positive cellular backgrounds, we selected representative genes for orthogonal validation by quantitative real-time PCR. Expression was examined in BJAB and BJAB-KSHV cells and across additional KSHV-negative and naturally infected B-cell models, including Ramos and BCBL1 cells and DG75 and BC3 cells. In parallel, we examined primary infection by exposing peripheral blood mononuclear cells (PBMCs) to purified KSHV (Fig. 3D). Analysis of selected kinetochore-associated transcripts across these models showed partial concordance with the expression patterns identified by transcriptomic analysis (Fig. 3E). CENPB and LIC2 were retained in this analysis as functionally important comparators despite their limited transcriptional response in the RNA-seq dataset.

We next asked whether individual KSHV proteins could contribute to the observed transcriptional changes. HEK293T cells were transfected with expression constructs encoding representative latent and lytic KSHV proteins, including LANA, RTA, vCyclin, vFLIP, and vGPCR. Expression of the respective viral proteins was confirmed by immunoblotting (Fig. 3F), after which transcript levels of the selected kinetochore-associated genes were quantified by real-time PCR. Expression of individual viral proteins produced gene-specific changes in kinetochore-associated transcripts (Fig. 3G). These observations suggest that the kinetochore-associated transcriptional phenotype observed in KSHV-positive cells may reflect the combined activities of multiple viral gene products rather than the action of a single viral regulator.

Together, these findings demonstrate that KSHV-positive cells exhibit selective transcriptional remodeling of kinetochore and chromosome segregation-associated genes. Importantly, transcriptional regulation and functional dependency were not necessarily coupled, as exemplified by CENP-B and LIC2, which were required for efficient episome maintenance despite showing little or no significant transcriptional alteration in the BJAB-KSHV RNA-seq analysis. These results distinguish two complementary aspects of the relationship between KSHV and the kinetochore machinery, KSHV infection-associated remodeling of selected host kinetochore pathways and a functional dependence on specific kinetochore-associated proteins for viral episome persistence.

### LANA associates with CENP-B and LIC2 and engages kinetochore-associated machinery during the cell cycle

Stable maintenance of KSHV episomes requires their coordinated replication and segregation during successive cell divisions. LANA provides the principal viral interface linking KSHV terminal repeat DNA to host chromatin and therefore represents a potential mechanism through which cellular chromosome-segregation machinery could contribute to viral genome persistence. Since CENP-B and LIC2 emerged as host factors required for efficient KSHV episome maintenance, we next investigated whether these proteins are associated with LANA.

We first examined the association of LANA with CENP-B and LIC by co-immunoprecipitation. Expression of FLAG-tagged LANA was confirmed by immunoblotting (Fig. 4A), and FLAG immunoprecipitation followed by immunoblot analysis demonstrated co-precipitation of CENP-B and LIC with LANA (Fig. 4B,C). These biochemical data indicate that CENP-B and LIC are present in LANA-containing protein complexes and provide a potential molecular link between these kinetochore-associated factors and KSHV episome maintenance.

**Figure 4.**
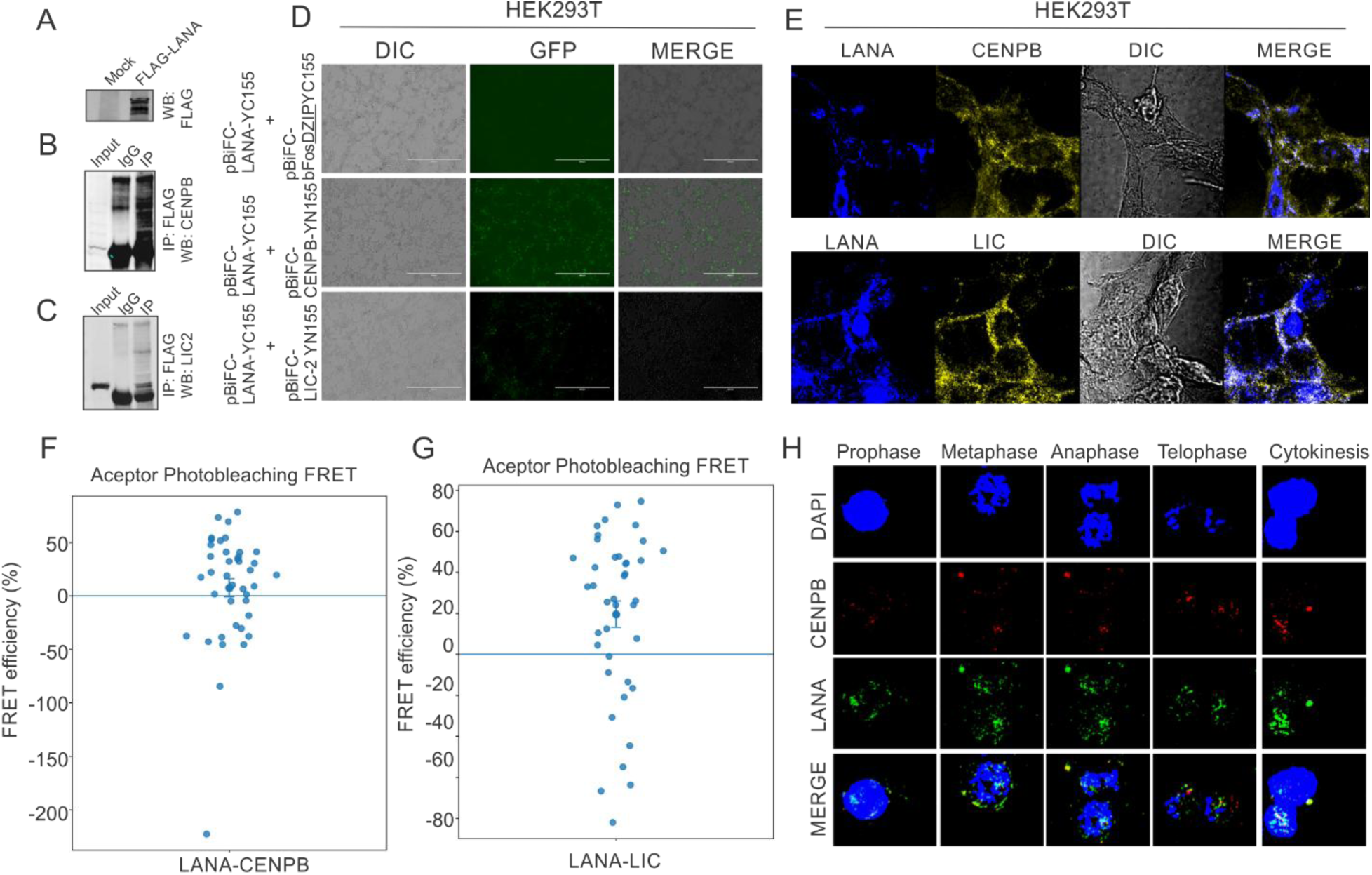
LANA associates with the kinetochore-associated proteins CENP-B and LIC2 and displays cell-cycle-dependent spatial association with kinetochore machinery. (A-C) Co-immunoprecipitation analysis of the association of LANA with CENP-B and LIC. Expression of FLAG-tagged LANA was confirmed by immunoblotting (A). FLAG immunoprecipitation followed by immunoblot analysis demonstrated co-precipitation of CENP-B (B) and LIC (C) with LANA. Input lysates and IgG controls are shown as indicated. (D) Bimolecular fluorescence complementation (BiFC) analysis of LANA association with CENP-B and LIC in intact cells. LANA and the indicated candidate proteins were fused to complementary fragments of the fluorescent reporter, and fluorescence reconstitution was monitored by microscopy. Representative DIC, fluorescence, and merged images of the indicated experimental and control combinations are shown. Reconstitution of fluorescence upon co-expression of the corresponding LANA and CENPB or LIC2 fusion constructs. (E) Fluorescence microscopy analysis of the intracellular distribution of LANA with CENP-B or LIC. Representative images show LANA, the corresponding kinetochore-associated protein, DIC, and merged channels. Overlapping fluorescence in the merged images indicates regions of spatial co-localization between LANA and CENPB or LIC2. (F, G) Acceptor-photobleaching FRET analysis of molecular proximity between LANA and CENPB (F) or LANA and LIC2 (G). FRET efficiency was determined from changes in donor fluorescence following photobleaching of the acceptor fluorophore. Individual points represent measurements from individual analyzed cells. (H) Cell-cycle-dependent localization of LANA relative to CENP-B during mitotic progression. Representative fluorescence images show cells at prophase, metaphase, anaphase, telophase, and cytokinesis. DNA was visualized with DAPI, and the corresponding DIC, CENP-B, LANA, and merged images are shown. The changing spatial distribution of LANA and CENP-B across successive stages of cell division illustrates their dynamic association during mitotic progression.

To determine whether LANA associates with these proteins in intact cells, we next employed bimolecular fluorescence complementation (BiFC). LANA and the candidate proteins were fused to complementary non-fluorescent fragments of the fluorescent reporter. Co-expression of the corresponding LANA and CENP-B or LIC2 fusion constructs resulted in reconstitution of partial and selective fluorescence, whereas the appropriate control combinations exhibited little or substantially lower fluorescence (Fig. 4D). These observations provide independent evidence that LANA and CENP-B/LIC2 can associate within the cellular environment.

We further investigated the intracellular distribution of these proteins by fluorescence microscopy followed by FRET based experiments. LANA exhibited overlapping spatial localization with both CENP-B and LIC in cells expressing the respective proteins (Fig. 4E). Although co-localization alone does not establish direct molecular interaction, these observations demonstrate that LANA and the candidate kinetochore-associated proteins occupy overlapping cellular regions and are consistent with the associations detected by co-immunoprecipitation and BiFC.

To examine molecular proximity, we performed acceptor-photobleaching fluorescence resonance energy transfer (FRET) analysis. LANA and CENP-B or LIC2 were expressed as donor- and acceptor-tagged fusion proteins, and FRET efficiency was determined from the increase in donor fluorescence following acceptor photobleaching. Individual-cell measurements revealed detectable FRET between LANA and CENP-B and between LANA and LIC (Fig. 4F,G), supporting close molecular proximity between the respective protein pairs. Together with the biochemical and BiFC analyses, these results provide complementary evidence for association of LANA with CENP-B and LIC2.

Because CENP-B and LIC2 participate in chromosome-associated processes relevant to episome inheritance, we next examined their spatial relationship with LANA during cell division. Imaging across successive stages of mitosis revealed dynamic localization of LANA relative to CENP-B, with spatial association evident during prophase, metaphase, anaphase, telophase, and cytokinesis (Fig. 4H). The distribution of the signals changed as chromosomes condensed, aligned, segregated, and entered daughter nuclei, indicating that the spatial relationship between LANA and kinetochore-associated machinery is cell-cycle dependent rather than static.

Collectively, co-immunoprecipitation, BiFC, fluorescence localization, FRET, and cell-cycle imaging provide complementary evidence that LANA associates closely with CENP-B and LIC2. These observations, together with the requirement for CENP-B and LIC2 in maintaining KSHV genome abundance, support a model in which LANA engages kinetochore-associated host factors as part of the machinery contributing to stable inheritance of viral episomes during cell division.

### Depletion of CENP-B and LIC disrupts chromosome segregation and promotes abnormal DNA content in KSHV-positive cells

Based on the observations that CENP-B and LIC2 contribute to KSHV episome maintenance and associate with LANA, we next investigated whether depletion of these proteins perturbs the host chromosome-segregation machinery. Because kinetochore function is required for accurate chromosome alignment and segregation during mitosis, we hypothesized that loss of CENP-B or LIC2 could produce mitotic abnormalities that accompany the defects in viral episome maintenance observed in the knockdown cells.

We first examined cellular DNA-content profiles by flow cytometry following propidium iodide staining. shControl cells exhibited the expected distribution of cells with predominant 2N and 4N DNA content, corresponding to G1 and G2/M populations, respectively (Fig. 5A,B). In contrast, depletion of CENP-B or LIC2 altered the DNA-content distribution and was accompanied by an increased population of cells containing DNA content greater than 4N (Fig. 5A,B). Quantification of the cell-cycle profiles demonstrated redistribution among the major DNA-content populations following depletion of either protein, with an enrichment of the >4N population compared with shControl cells (Fig. 5B). These findings indicate that depletion of CENP-B or LIC2 promotes accumulation of cells with abnormal DNA content consistent with defects in mitotic progression and/or chromosome segregation.

**Figure 5.**
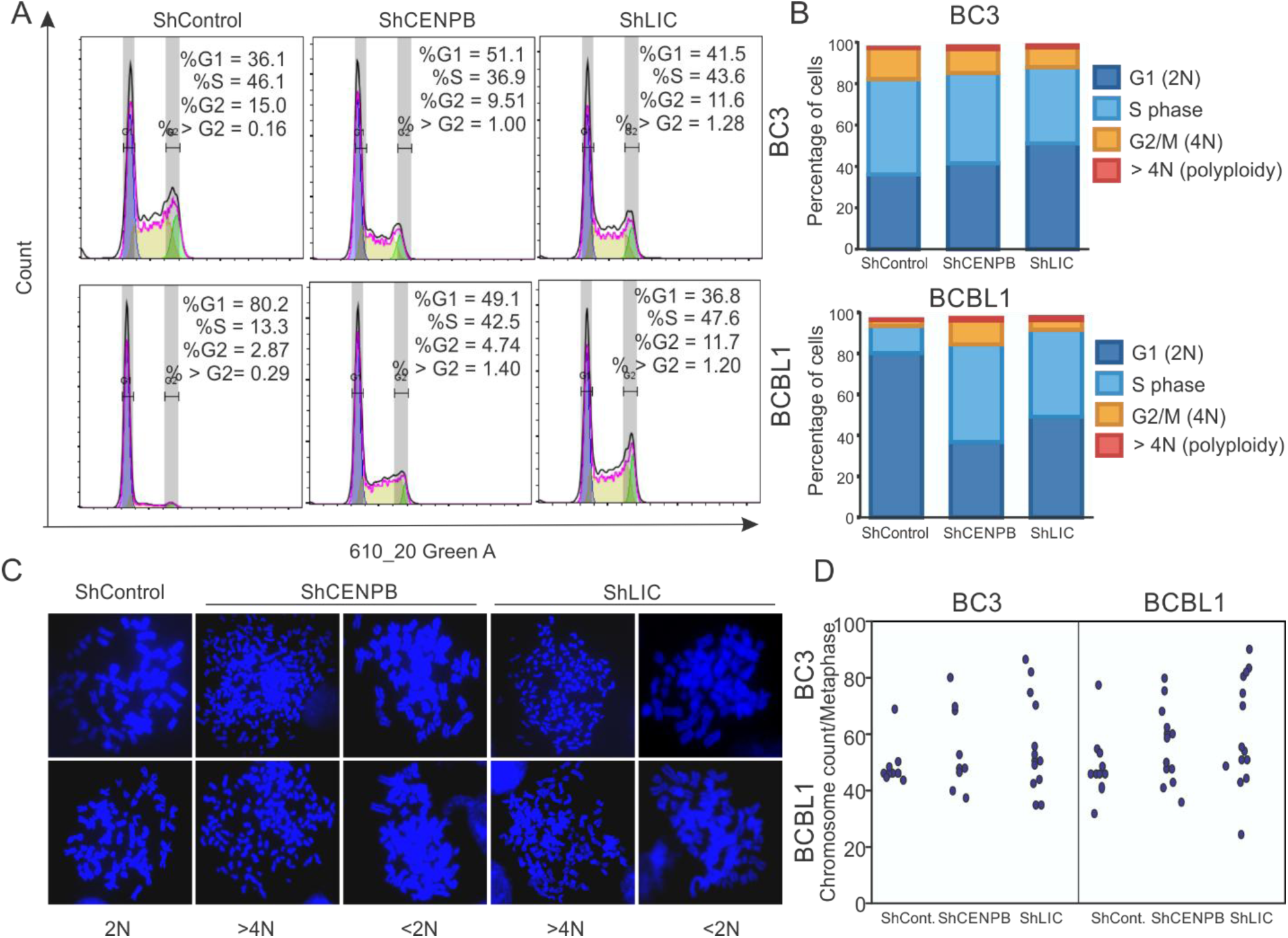
Depletion of CENPB and LIC2 disrupts chromosome segregation and promotes abnormal DNA content in KSHV-positive cells. (A) Representative cell-cycle DNA-content profiles of shControl-, shCENPB-, and shLIC2-expressing KSHV-positive cells analyzed by flow cytometry following propidium iodide (PI) staining. The positions of populations corresponding to approximately 2N, 4N, and >4N DNA content are indicated. (B) Quantification of DNA-content distribution in shControl, shCENPB, and shLIC2 cells. The relative proportions of cells within the G1/2N, S-phase, G2/M/4N, and >4N DNA-content populations are shown. (C) Representative images for mitotic chromosomes and nuclear abnormalities observed following CENPB or LIC2 depletion, including normal metaphase organization in shControl cells, polyploidy/aneuploidy in shCENPB-, and shLIC2 cells. (D) Quantification of mitotic chromosomes number in CENPB or LIC2 knock-down BC3 and BCBL1 cells.

To directly examine chromosome organization during mitosis, we next analyzed metaphase chromosome profiling by fluorescence microscopy. Representative examples of the major mitotic abnormalities observed following depletion of CENP-B or LIC2 including polyploidy or aneuploidy are shown in Fig. 5C. The quantification of cells with major mitotic abnormalities indicated significant effect on chromosome numbers in CENP-B or LIC2 knock-down cells

Together, these results demonstrate that depletion of CENP-B or LIC2 compromises faithful host chromosome organization and segregation during mitosis. The combination of metaphase misalignment, anaphase segregation defects, and accumulation of cells with >4N DNA content supports a role for these proteins in maintaining mitotic fidelity in KSHV-positive cells. In conjunction with the reduced KSHV genome abundance observed following CENP-B or LIC2 depletion, these findings support a model in which disruption of kinetochore-associated machinery simultaneously compromises host chromosome segregation and the stable inheritance of KSHV episomes (Fig. 5C,D).

### Kinetochore protein depletion restricts KSHV replication and impairs LANA–PCNA loading at viral episomes

As KSHV episome persistence requires replication of the viral genome in addition to its faithful segregation during cell division, we next investigated whether CENP-B and LIC2 contribute to KSHV DNA replication.

To examine viral DNA replication at the single-molecule level, we performed single-molecule analysis of replicated DNA (SMARD) in KSHV-positive BC3 cells in stable ShCENP-B or ShLIC2 cells as compared to ShControl cells (Fig. 6A). Sequential incorporation of nucleotide analogs into newly synthesized DNA was used to visualize replication patterns along individual DNA molecules. Representative replication tracks revealed differences between shControl cells and cells depleted of CENP-B or LIC (Fig. 6B). Quantitative analysis indicated reduced replication initiation and altered replication progression following depletion of both CENPB and LIC2 (Fig. 6B,C). These observations support a role for CENP-B and LIC in efficient replication of KSHV episomal DNA.

**Figure 6.**
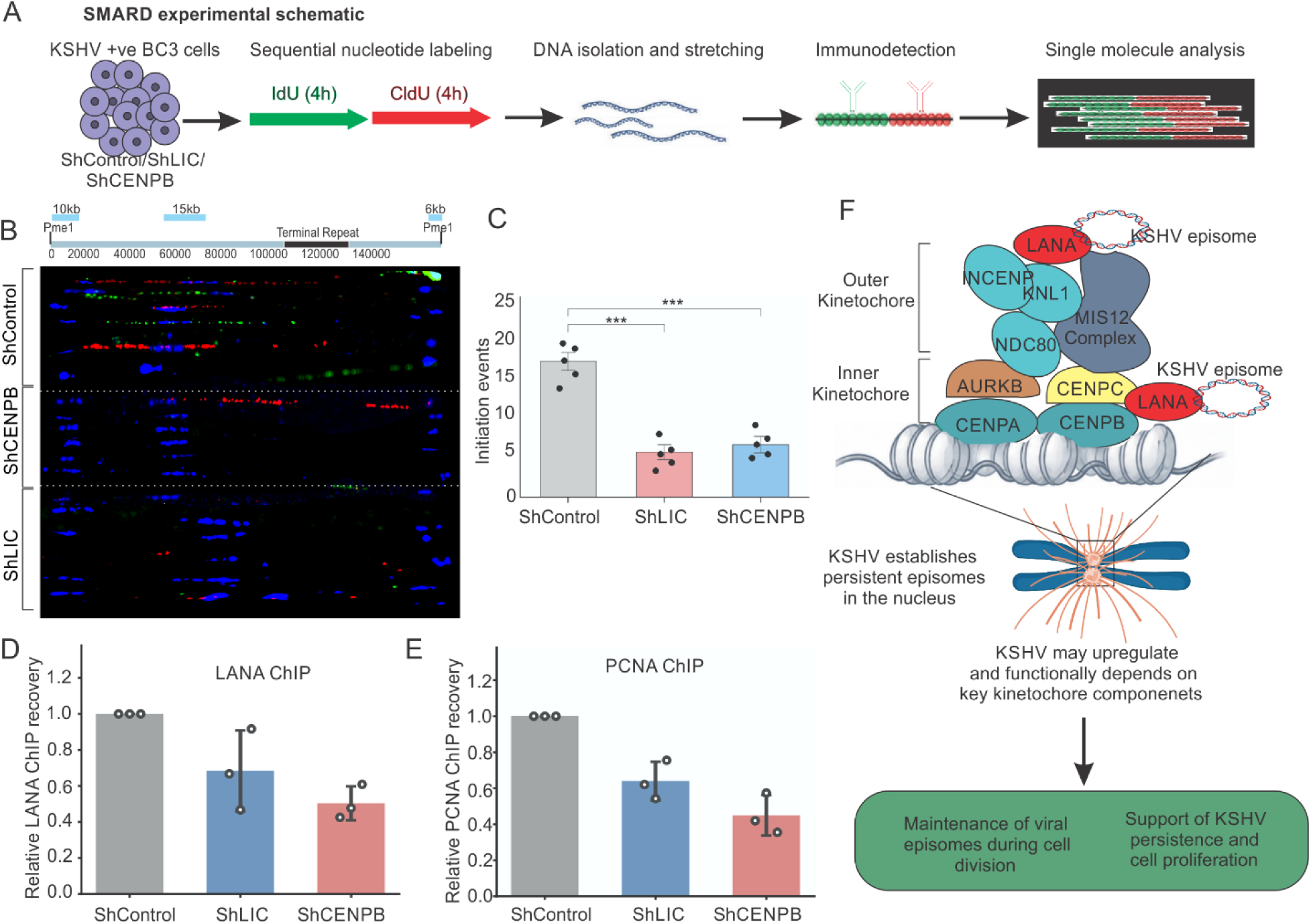
CENP-B and LIC2 depletion restricts KSHV episome replication and impairs LANA-PCNA recruitment to viral terminal repeats. **(A)** Schematic representation of the single-molecule analysis of replicated DNA (SMARD) workflow used to examine KSHV DNA replication following depletion of CENP-B or LIC2. KSHV-positive BC3 cells expressing control shControl, shCENPB, or shLIC2 were sequentially labeled with IdU and CldU, followed by isolation and stretching of genomic DNA and immunofluorescence detection of newly synthesized DNA molecules. **(B)** Representative single-molecule replication patterns from shControl, shCENPB, and shLIC2 cells. Sequential incorporation of IdU and CldU was visualized along individual DNA molecules to assess replication initiation and progression. Representative molecules from each condition are shown. **(C)** Quantification of replication initiation events in shControl, shCENPB, and shLIC2 cells. Initiation events were identified from the sequential nucleotide-labeling patterns and quantified across analyzed DNA molecules. **(D,E)** Chromatin immunoprecipitation followed by quantitative PCR (ChIP-qPCR) analysis of LANA and PCNA occupancy at the KSHV terminal repeat (TR) region following depletion of CENP-B or LIC2. Chromatin from shControl, shCENPB, and shLIC2 cells was immunoprecipitated using antibodies against LANA or PCNA, and enrichment of TR DNA was determined by qPCR. **(F)** Proposed model for the contribution of CENP-B and LIC to KSHV episome replication. The data supports a model in which CENPB and LIC2 promote efficient LANA occupancy at KSHV TR sequences and facilitates recruitment of PCNA and the host DNA replication machinery. Depletion of either reduces LANA and PCNA association with TR DNA and is accompanied by impaired viral DNA replication, thereby compromising efficient episome duplication and long-term persistence.

We next investigated a potential mechanism underlying the replication phenotype. LANA binds the KSHV terminal repeat (TR) sequences and provides an essential platform for episome maintenance and replication. We therefore examined whether depletion of CENP-B or LIC2 affects LANA occupancy at viral TR DNA. Chromatin immunoprecipitation followed by quantitative PCR (ChIP-qPCR) showed reduced enrichment of LANA at the TR following depletion of either CENP-B or LIC2 (Fig. 6D), suggesting that these kinetochore-associated proteins contribute to efficient LANA association with the viral episome.

Furthermore, as proliferating cell nuclear antigen (PCNA) functions as a processivity factor during DNA synthesis, we also examined its association with KSHV TR DNA. PCNA ChIP-qPCR similarly showed reduced enrichment at the TR following depletion of CENP-B or LIC2 (Fig. 6E). The parallel reduction in LANA and PCNA occupancy, together with the replication phenotype detected by SMARD, supports a mechanistic relationship between kinetochore-associated protein function, LANA occupancy, recruitment of cellular replication machinery, and efficient viral DNA synthesis.

Together, these results support a model in which CENP-B and LIC2 facilitate KSHV episome replication by promoting or stabilizing LANA association with viral TR sequences and supporting recruitment or retention of PCNA at the viral genome (Fig. 6F). Thus, in addition to contributing to chromosome-associated episome inheritance, CENP-B and LIC2 appear to support efficient duplication of KSHV episomal DNA, providing a potential mechanistic link between host kinetochore-associated machinery and the long-term persistence of KSHV genomes.

## Discussion

The faithful maintenance of Kaposi’s sarcoma-associated herpesvirus (KSHV) episomes is central to the establishment of lifelong latency and the development of KSHV-associated malignancies[5, 51, 52]. Although KSHV-encoded LANA has long been recognized as the principal viral factor anchoring episomes to host chromatin, growing evidence suggests that this process depends on host chromosome segregation machinery[5, 17, 53]. In this study, we systematically interrogated the contribution of 37 kinetochore-associated proteins to KSHV persistence and identified new host factors, including CENP-B and the Light Intermediate Chain (LIC2) of the dynein complex, as critical regulators of viral episome stability. These findings extend the repertoire of host proteins involved in γ-herpesvirus latency and highlight the broader exploitation of the kinetochore machinery by KSHV[54–56].

Our initial screen demonstrated that multiple kinetochore proteins are required for efficient maintenance of KSHV episomes. While previous studies had implicated CENP-F, Bub1, BubR1, Aurora B, Survivin, and NuMA in episome persistence, our results expand this list to include additional components, most notably CENP-B and LIC2. CENP-B is a DNA-binding protein that recognizes CENP-B boxes within centromeric DNA and is essential for centromere function[57–59]. Its newly identified interaction with LANA suggests that KSHV may utilize CENP-B to strengthen viral episome tethering at centromeres. Similarly, LIC2, a dynein subunit critical for microtubule-based transport and chromosome movement[60, 61], was shown to influence both episome maintenance and chromosomal segregation. The loss of LIC2 not only destabilized viral episomes but also promoted host chromosomal missegregation and polyploidy, suggesting that viral and cellular genomes are interdependent on shared kinetochore-dynein pathways.

Beyond identifying novel host factors, our study revealed that KSHV infection itself may actively influence kinetochore biology. We observed altered expressions of several kinetochore-encoding genes in KSHV-infected cells[50], suggesting that the virus may modulate host transcriptional networks to create a more favorable environment for episome persistence. This is consistent with prior observations that herpesviruses reprogram host epigenetic and signaling pathways to maintain latency[62]. Together, these findings imply a bidirectional relationship in which KSHV both depends on and reshapes kinetochore pathways to optimize episome maintenance.

Our validation experiments in naturally infected KSHV-positive cell lines strengthen the physiological relevance of these findings. Unlike transient models, naturally infected cells preserve the native interplay between viral and host proteins, providing confidence that kinetochore factors, including CENP-B and LIC2, contribute to episome persistence under biologically relevant conditions. Furthermore, the SMARD experiments offered mechanistic insight by demonstrating that knockdown of CENP-B or LIC2 restricts replication initiation events, resulting in reduced episome duplication. This indicates that kinetochore proteins not only stabilize episome segregation but also facilitate efficient viral DNA replication.

Taken together, our results support a model in which KSHV episome maintenance is safeguarded by a network of kinetochore-associated proteins. LANA anchors episomes at chromosomal sites, where interactions with kinetochore associate proteins such as CENP-B enhance centromere association and LIC2 integrates viral genomes into microtubule-based segregation machinery. Disruption of these interactions impairs viral episome stability, persistence and host chromosome fidelity, highlighting the integration of KSHV into the cell division apparatus.

In conclusion, this study provides a systematic analysis of kinetochore-associated proteins in KSHV latency, revealing CENP-B and LIC2 as novel host dependency factors. These findings broaden our understanding of host–virus interactions at the chromosome segregation interface and suggest new avenues for therapeutic intervention. Targeting kinetochore-virus interactions could represent a strategy to destabilize viral episomes and eliminate latent reservoirs, a central challenge in the treatment of KSHV-associated malignancies. Future studies will be necessary to define the structural basis of these interactions and to elucidate how dynein subunits mediate episome partitioning and determine whether similar mechanisms are exploited by other oncogenic Ƴ-herpesviruses.

## Materials and Methods

### Cell Culture and Maintenance

BJAB, BJAB-KSHV, BC3, BCBL1, Ramos, and DG75 cells were maintained in RPMI 1640 medium supplemented with 7% bovine growth serum (BGS) and penicillin/streptomycin at 37 °C in a humidified atmosphere containing 5% CO₂. BJAB-KSHV cells were cultured in the presence of 2 µg/mL puromycin to maintain selection of the KSHV-positive population.

Human embryonic kidney HEK293T cells and HEK293T cells harboring recombinant BAC16-KSHV (HEK293T-BAC16-KSHV) were maintained in DMEM supplemented with 7% BGS and penicillin/streptomycin at 37 °C in a humidified atmosphere containing 5% CO₂. HEK293T-BAC16-KSHV cells were additionally maintained under selection with 200 µg/mL hygromycin. Peripheral blood mononuclear cells (PBMCs) from de-identified healthy donors were obtained through the Human Immunology Core (HIC) at the University of Pennsylvania. Donor samples were collected under Institutional Review Board (IRB) approved protocols administered by the HIC, in accordance with the principles of the Declaration of Helsinki. Written informed consent was obtained from all donors in accordance with the approved protocols.

### KSHV Virion Preparation and Infection of Peripheral Blood Mononuclear Cells

KSHV virions were produced from iSLK-BAC-KSHV cells following induction of the viral lytic cycle. Cells were treated with 12-O-tetradecanoylphorbol-13-acetate (TPA; 20 ng/mL) and sodium butyrate (3 mM) and maintained at 37 °C in 5% CO₂ for 5 days. At the end of the induction period, cells and culture supernatants were separated by centrifugation at 3,000 rpm for 30 min. The clarified culture supernatant was passed through a 0.45-µm filter.

The harvested cell pellet was resuspended in 10 mL of 1× phosphate-buffered saline (PBS) and subjected to three freeze–thaw cycles to release cell-associated viral particles. Cellular debris was removed by centrifugation at 3,000 rpm for 30 min, and the resulting supernatant was passed through a 0.45-µm filter. The filtrates obtained from the culture medium and freeze–thaw lysate were pooled, and KSHV virions were concentrated by ultracentrifugation at 23,500 rpm for 2 h. The resulting viral preparation was used for infection of PBMCs in the presence of 8 µg/mL polybrene at a MOI of 1:10.

### Plasmid Constructs and Transfection

Expression constructs encoding the KSHV proteins LANA, RTA, vCyclin, vFLIP, and vGPCR were generated in the pLVX-AC-GFP expression vector, producing GFP-tagged fusion proteins. HEK293T cells were transiently transfected with the indicated plasmid expression using jetPRIME transfection reagent (Polyplus) according to the manufacturer’s instructions. Cells transfected with the parental pLVX-AC-GFP vector were used as controls. Expression of the GFP-tagged viral proteins was confirmed by immunoblotting prior to downstream gene-expression analysis. For co-immunoprecipitation experiments, a FLAG-tagged LANA expression construct (pA3F-LANA) was used to examine the association of LANA with endogenous CENP-B and LIC.

For bimolecular fluorescence complementation (BiFC) analysis, LANA was cloned into the pBiFC-bFosYC155 vector, whereas CENP-B and LIC were individually cloned into the pBiFC-bJunYN155 vector. The resulting LANA-YC155 and CENP-B-YN155 or LIC-YN155 fusion constructs were co-expressed in HEK293T cells. Reconstitution of the complementary fluorescent-protein fragments was subsequently evaluated by fluorescence microscopy

For fluorescence resonance energy transfer (FRET) experiments, LANA was expressed as a cyan fluorescent protein (CFP) fusion using the pECFP-C1 vector, whereas CENP-B and LIC were expressed as yellow fluorescent protein (YFP) fusion proteins using the pEYFP-C1 vector. The resulting CFP-LANA and YFP-CENP-B or YFP-LIC constructs were co-expressed in HEK293T cells for analysis of molecular proximity by acceptor-photobleaching FRET.

### shRNA Constructs, Lentiviral Production, and Stable Knockdown

Short hairpin RNA (shRNA) constructs targeting kinetochore- and chromosome segregation-associated genes were used to systematically evaluate their contribution to KSHV genome maintenance. Two independent shRNA constructs targeting each candidate gene were initially evaluated for knockdown efficiency in HEK293T cells. HEK293T cells were transfected with the individual shRNA expression constructs using jetPRIME transfection reagent (Polyplus), and target-gene expression was subsequently measured by quantitative reverse-transcription PCR (qRT-PCR). For each candidate gene, the shRNA producing the greater reduction in transcript abundance was selected for subsequent functional analysis. A non-targeting shRNA construct (shControl) was used as reference control throughout the experiments.

For the initial functional screen, the selected shRNA construct for each candidate gene was transfected into HEK293T-BAC16-KSHV cells using jetPRIME transfection reagent. KSHV genome abundance following target-gene depletion was subsequently determined by quantitative PCR as described below.

For stable depletion of CENP-B and LIC in BC3 and BCBL1 cells, shRNA constructs were delivered by lentiviral transduction. Lentiviral particles were produced in HEK293T cells by co-transfection of the individual shRNA expression constructs together with lentiviral packaging plasmids. Virus-containing culture supernatants were harvested and clarified by centrifugation at 3,000 rpm for 30 min to remove cellular debris, followed by passage through a 0.45-µm filter. Lentiviral particles were concentrated from the clarified supernatants by ultracentrifugation at 23,500 rpm for 2 h. BC3 and BCBL1 cells were transduced with the concentrated lentiviral preparations in the presence of 8 µg/mL polybrene. Transduced cells were selected with 2 µg/mL puromycin, and the resulting stable knockdown populations were subsequently maintained in medium containing 1 µg/mL puromycin. Depletion of CENP-B and LIC was confirmed by qRT-PCR using RNA isolated from independent cell preparations prior to their use in subsequent experiments. Primer sequences for qRT-PCR and Target sequences for all shRNA constructs are provided in Table S1 and Table S2 respectively.

### RNA Isolation, cDNA Synthesis, and Quantitative Real-Time PCR

Total RNA was isolated from cultured cells using TRIzol reagent (Ambion) according to the manufacturer’s instructions. Following chloroform-mediated phase separation, RNA was recovered from the aqueous phase by precipitation, washed, and resuspended in RNase-free water. RNA concentration and purity were determined prior to cDNA synthesis.

For reverse transcription, 2 µg of total RNA was used for first-strand cDNA synthesis using a SuperScript cDNA synthesis kit (Applied Biosystems) with random primers according to the manufacturer’s instructions. The resulting cDNA was diluted 10-fold with nuclease-free water prior to quantitative real-time PCR (qRT-PCR).

Quantitative PCR was performed using SYBR Green PCR Master Mix (Applied Biosystems) on a QuantStudio real-time PCR system. One microliter of diluted cDNA was used as template for each reaction. Gene-specific primer pairs were used to quantify transcript abundance. Relative gene expression was calculated using the comparative delta delta CT method using GAPDH as an endogenous control. For Absolute quantification of KSHV genome, KSHV genome copy number was determined by absolute quantitative PCR using a plasmid-based standard-curve method as described previously[63]. Primer sequences used for KSHV genome quantification are provided in Table S1.

### Immunoblotting and Co-immunoprecipitation

Whole-cell lysates were prepared in radioimmunoprecipitation assay (RIPA) buffer containing 50 mM Tris-HCl (pH 7.6), 150 mM NaCl, 2 mM EDTA, and 1% Nonidet P-40, supplemented with the protease inhibitors aprotinin, leupeptin, pepstatin, and 1 mM phenylmethylsulfonyl fluoride (PMSF). Protein concentrations were determined, and equal amounts of total protein were resolved by sodium dodecyl sulfate-polyacrylamide gel electrophoresis (SDS-PAGE) and transferred onto nitrocellulose membranes. Membranes were blocked with 5% nonfat dry milk prepared in Tris-buffered saline containing Tween 20 (TBST) for 1 h at room temperature and subsequently incubated with the indicated primary antibodies. Following incubation with appropriate infrared dye-conjugated secondary antibodies, immunoreactive bands were visualized using an Odyssey infrared imaging system (LI-COR Biosciences). GAPDH was used as a loading control where indicated.

For co-immunoprecipitation experiments, HEK293T cells transfected with FLAG-LANA (pA3F-LANA) or the corresponding control vector were lysed in RIPA buffer containing protease inhibitors. Equal amounts of protein lysate were pre-cleared with protein A/G agarose beads and subsequently incubated with anti-FLAG antibody overnight at 4 °C with gentle mixing. Immune complexes were captured using protein A/G agarose beads and washed three times with 1× phosphate-buffered saline (PBS). Bound proteins were eluted by resuspending the beads in 2× SDS sample buffer and analyzed by SDS-PAGE and immunoblotting. Approximately 5% of the lysate used for immunoprecipitation was retained as the input control. Immunoprecipitated FLAG-LANA and associated endogenous CENP-B and LIC were detected using the corresponding antibodies.

### Immunofluorescence Microscopy and LANA Nuclear-Foci Quantification

For immunofluorescence analysis, cells were collected, washed with phosphate-buffered saline (PBS), and semi-dried on 8-well glass slides. Cells were fixed with (4% paraformaldehyde). Single step permeabilized and blocking was performed using 0.3% Triton-X100 in PBS supplemented with 5% normal serum. Samples were subsequently incubated with an anti-LANA primary antibody followed by an appropriate fluorophore-conjugated secondary antibody. Nuclei were counterstained with DAPI, and slides were mounted using antifade mounting medium.

Fluorescence images were acquired on Olympus IX81 confocal microscope. LANA nuclear foci were quantified from individual cells using ImageJ software. Nuclei were identified using DAPI staining, and discrete intranuclear LANA-positive foci were counted within individual nuclear regions of interest. Cells from multiple fields were analyzed for each condition, and the number of LANA foci per nucleus was used as a measure of episome-associated LANA nuclear puncta. The number of cells and independent experiments analyzed is indicated in the corresponding figure legend.

### Bimolecular Fluorescence Complementation and FRET Analysis

Bimolecular fluorescence complementation (BiFC) was used to examine the association of LANA with CENP-B and LIC in intact cells. LANA was expressed as a fusion with the YC155 fragment using the pBiFC-bFosYC155 vector, whereas CENP-B and LIC were individually expressed as fusions with the complementary YN155 fragment using the pBiFC-bJunYN155 vector. HEK293T cells were co-transfected with the indicated combinations of LANA-YC155 and CENP-B-YN155 or LIC-YN155 constructs using jetPRIME transfection reagent. BiFC fluorescence resulting from complementation of the YN155 and YC155 fragments was evaluated by fluorescence microscopy after 48h of transfection.

Fluorescence resonance energy transfer (FRET) was independently used to evaluate the close molecular proximity of LANA with CENP-B or LIC. HEK293T cells were co-transfected with CFP-LANA together with YFP-CENP-B or YFP-LIC expression constructs generated using the pECFP-C1 and pEYFP-C1 vectors, respectively. FRET was measured using the acceptor-photobleaching method. CFP donor and YFP acceptor fluorescence images were acquired before photobleaching, after which the YFP acceptor was selectively photobleached. CFP donor fluorescence was subsequently re-acquired using identical imaging conditions. FRET efficiency was determined from the increase in CFP donor fluorescence following acceptor photobleaching according to the formula, FRET efficiency (%) = [(D_post − D_pre)/D_post] × 100, where D_pre and D_post represent CFP donor fluorescence intensities before and after acceptor photobleaching, respectively. Individual cells were treated as measurement units for FRET quantification.

### Cell-Cycle Analysis and Chromosome-Segregation Imaging

Cell-cycle distribution and DNA content were analyzed by propidium iodide (PI) staining followed by flow cytometry. ShControl, ShCENPB or ShLIC cells were harvested, washed with PBS, fixed in 70% ethanol. Cells were stained with PI in the presence of RNase A. DNA-content profiles were acquired by flow cytometry and analyzed using FlowJo software. The proportions of cells within G0/G1, S, and G2/M phases, as well as populations exhibiting abnormal DNA content (>4N), were quantified using identical analysis criteria across experimental groups.

Chromosome-segregation phenotypes were evaluated by fluorescence microscopy of intact mitotic cells. Metaphase cells were assessed for chromosome alignment at the metaphase plate, whereas anaphase cells were examined for lagging chromosomes and other segregation abnormalities. Images were acquired using IX70 fluorescence microscope, and the frequencies of abnormal mitotic cells were quantified using the same scoring criteria across control and knockdown conditions.

### Single-Molecule Analysis of Replicated DNA (SMARD)

Single-molecule analysis of replicated DNA (SMARD) was performed using a modification of previously described methods[64, 65]. Cells were sequentially pulse-labeled with the thymidine analogs CldU (10 µM) and IdU (10 µM) and embedded in InCert agarose plugs. Following proteinase K digestion, genomic DNA was digested with PmeI to linearize the KSHV genome. DNA-containing agarose plugs were separated by pulsed-field gel electrophoresis (PFGE). Several gel slices corresponding approximately to the expected size of the linearized KSHV genome were collected and tested for enrichment of KSHV DNA. KSHV-enriched slices were identified by quantitative PCR using KSHV-specific primers. Selected agarose slices were subsequently digested with agarase to release the DNA, which was molecularly stretched onto silanized glass slides.

KSHV DNA molecules were identified using KSHV-specific hybridization probes, and incorporated CldU and IdU were detected using nucleotide analog-specific antibodies. Replication patterns were visualized by fluorescence microscopy, and individual KSHV DNA molecules were analyzed for replication initiation patterns and labeled replication-track lengths.

### Chromatin Immunoprecipitation and ChIP-qPCR

Chromatin immunoprecipitation (ChIP) was performed using a previously established protocol with minor modifications[62]. Cells were cross-linked with 1% formaldehyde at room temperature, and cross-linking was reversed with glycine. Cells were washed with PBS and lysed in cell lysis buffer (5 mM PIPES, pH 8.0, 85 mM KCl, and 0.5% NP-40) supplemented with protease inhibitors. Nuclei were isolated and lysed in nuclear lysis buffer (50 mM Tris-HCl, pH 8.0, 10 mM EDTA, and 1% SDS), and chromatin was fragmented to an average size of approximately 300–400 bp using a QSONICA sonicator with alternating 15-s sonication and 15-s cooling cycles for a total of 8 min.

Following centrifugation, soluble chromatin was diluted fivefold in ChIP-dilution buffer and pre-cleared with protein A/G agarose beads. Chromatin was incubated overnight at 4 °C with antibodies against LANA or PCNA, or with the corresponding control IgG. Immune complexes were captured with protein A/G agarose beads, washed, and eluted in 1% SDS/0.1 M NaHCO₃. Cross-links were reversed at 65 °C, followed by RNase and proteinase K treatment, and DNA was purified using a Qiagen column.

Purified ChIP DNA was analyzed by quantitative PCR using primers targeting the KSHV terminal repeat (TR) region. An aliquot of chromatin collected before immunoprecipitation was retained as DNA, and enrichment was calculated relative to input chromatin. Primer sequences used for ChIP-qPCR are provided in Table S1.

### Statistical Analysis

Quantitative data are presented as mean± standard deviation (SD), unless otherwise indicated. Statistical analyses were performed using GraphPad Prism. Comparisons between two groups were performed using two-tailed Student’s *t*-test, whereas experiments involving multiple groups were analyzed using one-way analysis of variance (ANOVA) followed by an appropriate multiple-comparison test, as indicated in the corresponding figure legends. A *P* value < 0.05 was considered statistically significant.

## Funding

This work was supported by the National Cancer Institute of the National Institutes of Health under Public Health Service grants P01-CA281867 and R01-CA268998 (to E.S.R.). The funders had no role in study design, data collection and interpretation, or the decision to submit the work for publication.

## Data Availability

All other data supporting the findings of this study are available within the article and its Supplementary Information files or from the corresponding author upon reasonable request.

---

**Supplementary Table 1:**
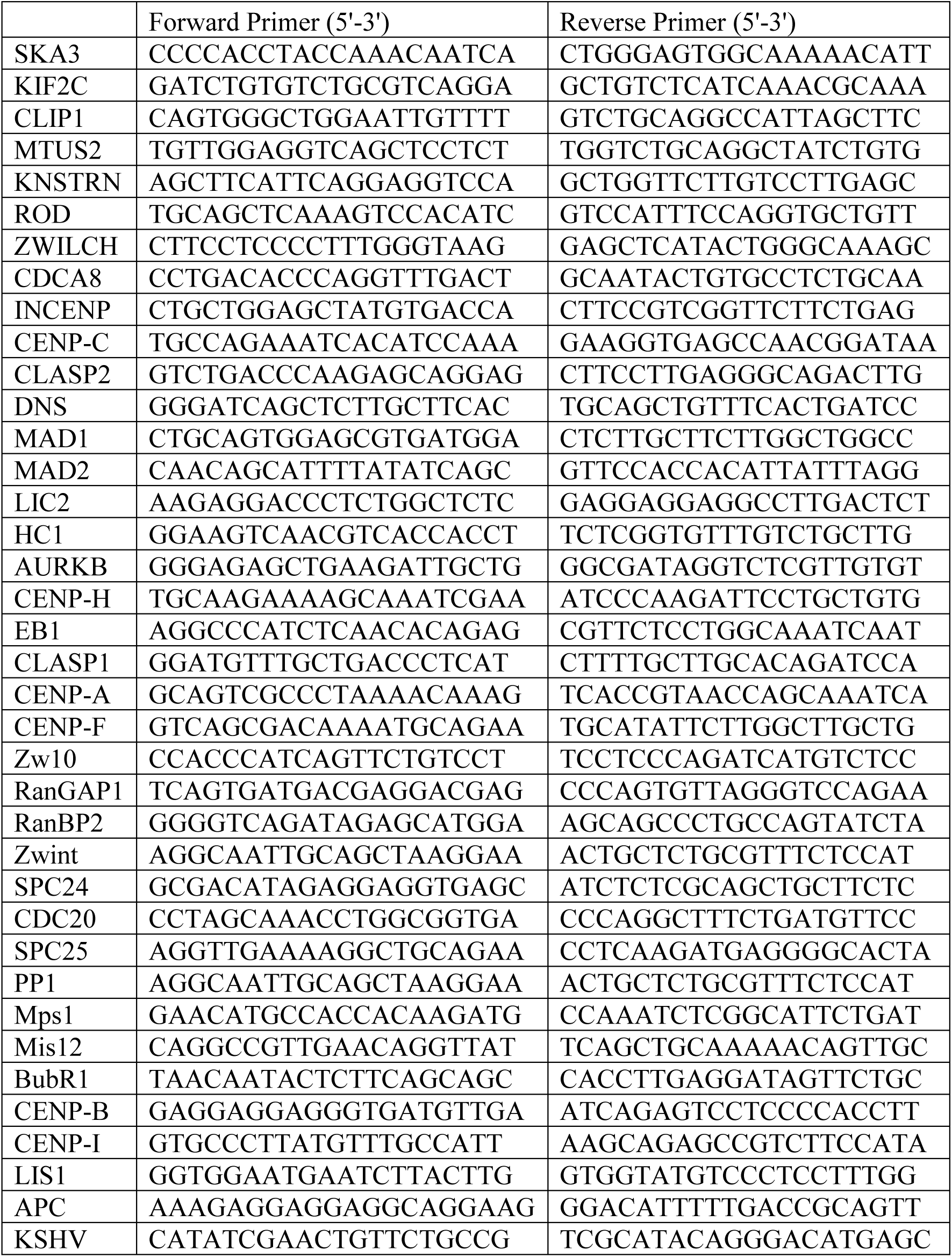
Sequences of qPCR primers used in the study.

**Supplementary Table 2:**
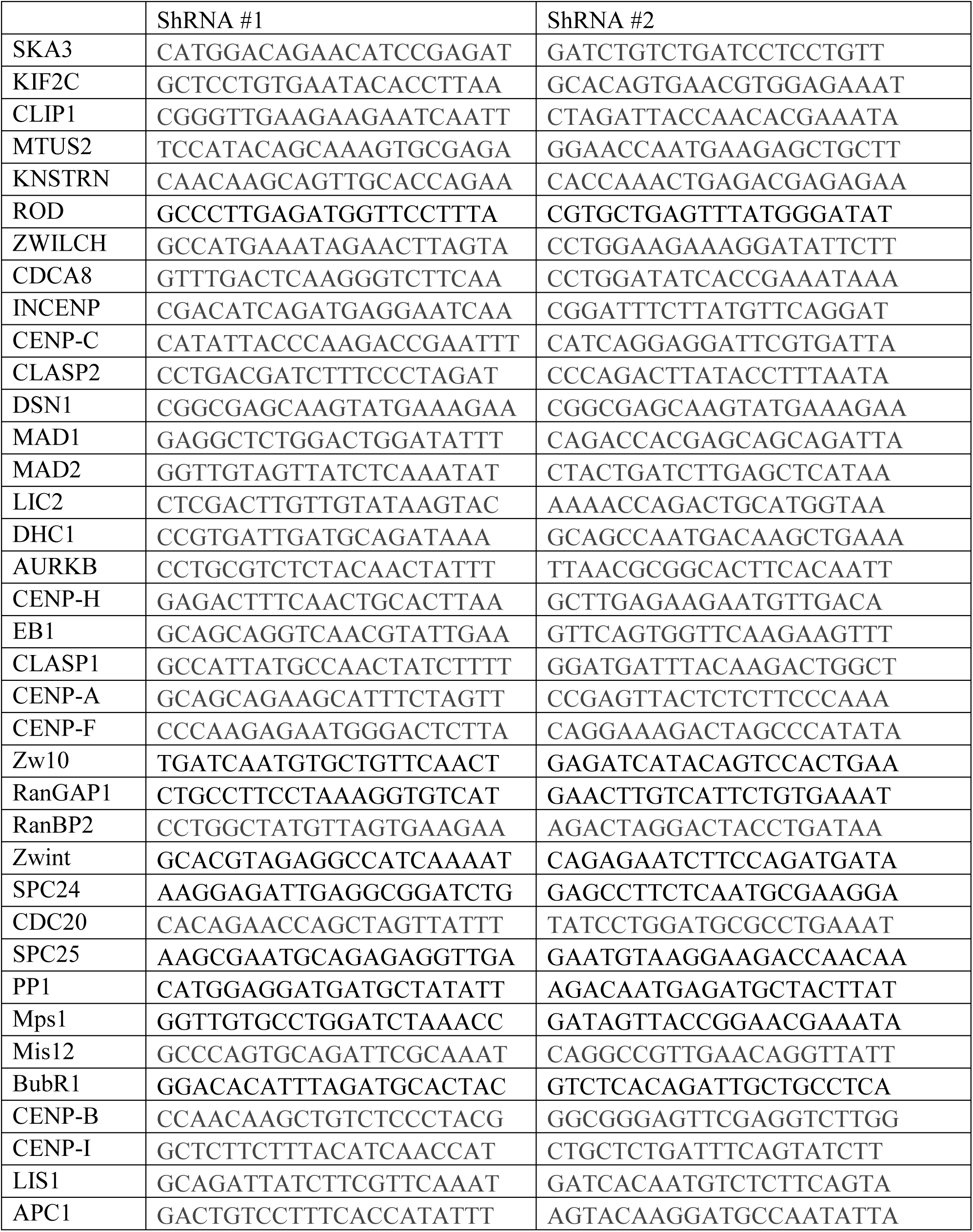
Sequence of ShRNA oligos.

## Notes

### Competing Interest Statement

The authors have declared no competing interest.

## References

1. Goncalves PH, Ziegelbauer J, Uldrick TS, Yarchoan R. Kaposi sarcoma herpesvirus-associated cancers and related diseases. Curr Opin HIV AIDS. 2017;12(1):47–56. doi: 10.1097/COH.0000000000000330. PubMed PMID: 27662501; PubMed Central PMCID: PMCPMC6311702.

2. Mesri EA, Cesarman E, Boshoff C. Kaposi’s sarcoma and its associated herpesvirus. Nat Rev Cancer. 2010;10(10):707–19. doi: 10.1038/nrc2888. PubMed PMID: 20865011; PubMed Central PMCID: PMCPMC4721662.

3. Cesarman E, Chang Y, Moore PS, Said JW, Knowles DM. Kaposi’s sarcoma-associated herpesvirus-like DNA sequences in AIDS-related body-cavity-based lymphomas. N Engl J Med. 1995;332(18):1186–91. doi: 10.1056/NEJM199505043321802. PubMed PMID: 7700311.

4. Ganem D. KSHV and the pathogenesis of Kaposi sarcoma: listening to human biology and medicine. J Clin Invest. 2010;120(4):939–49. doi: 10.1172/JCI40567. PubMed PMID: 20364091; PubMed Central PMCID: PMCPMC2847423.

5. Juillard F, Tan M, Li S, Kaye KM. Kaposi’s Sarcoma Herpesvirus Genome Persistence. Front Microbiol. 2016;7:1149. doi: 10.3389/fmicb.2016.01149. PubMed PMID: 27570517; PubMed Central PMCID: PMCPMC4982378.

6. Ye F, Lei X, Gao SJ. Mechanisms of Kaposi’s Sarcoma-Associated Herpesvirus Latency and Reactivation. Adv Virol. 2011;2011. doi: 10.1155/2011/193860. PubMed PMID: 21625290; PubMed Central PMCID: PMCPMC3103228.

7. Murdock SJ, Bersonda JR, Forrest JC, Manzano M. Molecular Mechanisms of KSHV Latency Establishment and Maintenance. Curr Clin Microbiol Rep. 2024;11(4):220–30. doi: 10.1007/s40588-024-00232-x. PubMed PMID: 40772265; PubMed Central PMCID: PMCPMC12327057.

8. Lieberman PM. Keeping it quiet: chromatin control of gammaherpesvirus latency. Nat Rev Microbiol. 2013;11(12):863–75. doi: 10.1038/nrmicro3135. PubMed PMID: 24192651; PubMed Central PMCID: PMCPMC4544771.

9. Ohsaki E, Ueda K. Kaposi’s Sarcoma-Associated Herpesvirus Genome Replication, Partitioning, and Maintenance in Latency. Front Microbiol. 2012;3:7. doi: 10.3389/fmicb.2012.00007. PubMed PMID: 22291692; PubMed Central PMCID: PMCPMC3264903.

10. Si H, Verma SC, Robertson ES. Proteomic analysis of the Kaposi’s sarcoma-associated herpesvirus terminal repeat element binding proteins. J Virol. 2006;80(18):9017–30. doi: 10.1128/JVI.00297-06. PubMed PMID: 16940514; PubMed Central PMCID: PMCPMC1563930.

11. Ballestas ME, Kaye KM. The latency-associated nuclear antigen, a multifunctional protein central to Kaposi’s sarcoma-associated herpesvirus latency. Future Microbiol. 2011;6(12):1399–413. doi: 10.2217/fmb.11.137. PubMed PMID: 22122438; PubMed Central PMCID: PMCPMC3857968.

12. Grundhoff A, Ganem D. The latency-associated nuclear antigen of Kaposi’s sarcoma-associated herpesvirus permits replication of terminal repeat-containing plasmids. J Virol. 2003;77(4):2779–83. doi: 10.1128/jvi.77.4.2779-2783.2003. PubMed PMID: 12552022; PubMed Central PMCID: PMCPMC141125.

13. Jain V, Renne R. Visualization of molecular biology: The LANA tether. Proc Natl Acad Sci U S A. 2018;115(19):4816–8. doi: 10.1073/pnas.1804797115. PubMed PMID: 29691321; PubMed Central PMCID: PMCPMC5949013.

14. Hellert J, Weidner-Glunde M, Krausze J, Lunsdorf H, Ritter C, Schulz TF, et al. The 3D structure of Kaposi sarcoma herpesvirus LANA C-terminal domain bound to DNA. Proc Natl Acad Sci U S A. 2015;112(21):6694–9. doi: 10.1073/pnas.1421804112. PubMed PMID: 25947153; PubMed Central PMCID: PMCPMC4450395.

15. Mercier A, Arias C, Madrid AS, Holdorf MM, Ganem D. Site-specific association with host and viral chromatin by Kaposi’s sarcoma-associated herpesvirus LANA and its reversal during lytic reactivation. J Virol. 2014;88(12):6762–77. doi: 10.1128/JVI.00268-14. PubMed PMID: 24696474; PubMed Central PMCID: PMCPMC4054346.

16. Kumar A, Lyu Y, Yanagihashi Y, Chantarasrivong C, Majerciak V, Salemi M, et al. KSHV episome tethering sites on host chromosomes and regulation of latency-lytic switch by CHD4. Cell Rep. 2022;39(6):110788. doi: 10.1016/j.celrep.2022.110788. PubMed PMID: 35545047; PubMed Central PMCID: PMCPMC9153692.

17. Uppal T, Banerjee S, Sun Z, Verma SC, Robertson ES. KSHV LANA--the master regulator of KSHV latency. Viruses. 2014;6(12):4961–98. doi: 10.3390/v6124961. PubMed PMID: 25514370; PubMed Central PMCID: PMCPMC4276939.

18. Xiao B, Verma SC, Cai Q, Kaul R, Lu J, Saha A, et al. Bub1 and CENP-F can contribute to Kaposi’s sarcoma-associated herpesvirus genome persistence by targeting LANA to kinetochores. J Virol. 2010;84(19):9718–32. doi: 10.1128/JVI.00713-10. PubMed PMID: 20660191; PubMed Central PMCID: PMCPMC2937805.

19. Verma SC, Lan K, Robertson E. Structure and function of latency-associated nuclear antigen. Curr Top Microbiol Immunol. 2007;312:101–36. doi: 10.1007/978-3-540-34344-8_4. PubMed PMID: 17089795; PubMed Central PMCID: PMCPMC3142369.

20. Gupta N, Thakker S, Verma SC. KSHV encoded LANA recruits Nucleosome Assembly Protein NAP1L1 for regulating viral DNA replication and transcription. Sci Rep. 2016;6:32633. doi: 10.1038/srep32633. PubMed PMID: 27599637; PubMed Central PMCID: PMCPMC5013526.

21. Sun Q, Tsurimoto T, Juillard F, Li L, Li S, De Leon Vazquez E, et al. Kaposi’s sarcoma-associated herpesvirus LANA recruits the DNA polymerase clamp loader to mediate efficient replication and virus persistence. Proc Natl Acad Sci U S A. 2014;111(32):11816–21. doi: 10.1073/pnas.1404219111. PubMed PMID: 25071216; PubMed Central PMCID: PMCPMC4136584.

22. Singh RK, Lamplugh ZL, Lang F, Yuan Y, Lieberman P, You J, et al. KSHV-encoded LANA protects the cellular replication machinery from hypoxia induced degradation. PLoS Pathog. 2019;15(9):e1008025. doi: 10.1371/journal.ppat.1008025. PubMed PMID: 31479497; PubMed Central PMCID: PMCPMC6743784.

23. Nakajima Ki, Inagaki T, Espera JM, Izumiya Y. Kaposi’s sarcoma-associated herpesvirus (KSHV) LANA prevents KSHV episomes from degradation. J Virol. 2024;98(2):e0126823. doi: 10.1128/jvi.01268-23. PubMed PMID: 38240588; PubMed Central PMCID: PMCPMC10878079.

24. Magidson V, O’Connell CB, Loncarek J, Paul R, Mogilner A, Khodjakov A. The spatial arrangement of chromosomes during prometaphase facilitates spindle assembly. Cell. 2011;146(4):555–67. doi: 10.1016/j.cell.2011.07.012. PubMed PMID: 21854981; PubMed Central PMCID: PMCPMC3291198.

25. Walczak CE, Cai S, Khodjakov A. Mechanisms of chromosome behaviour during mitosis. Nat Rev Mol Cell Biol. 2010;11(2):91–102. doi: 10.1038/nrm2832. PubMed PMID: 20068571; PubMed Central PMCID: PMCPMC2893392.

26. Purushothaman P, McDowell ME, McGuinness J, Salas R, Rumjahn SM, Verma SC. Kaposi’s sarcoma-associated herpesvirus-encoded LANA recruits topoisomerase IIbeta for latent DNA replication of the terminal repeats. J Virol. 2012;86(18):9983–94. doi: 10.1128/JVI.00839-12. PubMed PMID: 22761383; PubMed Central PMCID: PMCPMC3446556.

27. Rahayu R, Ohsaki E, Omori H, Ueda K. Localization of latency-associated nuclear antigen (LANA) on mitotic chromosomes. Virology. 2016;496:51–8. doi: 10.1016/j.virol.2016.05.020. PubMed PMID: 27254595.

28. Sun Z, Xiao B, Jha HC, Lu J, Banerjee S, Robertson ES. Kaposi’s sarcoma-associated herpesvirus-encoded LANA can induce chromosomal instability through targeted degradation of the mitotic checkpoint kinase Bub1. J Virol. 2014;88(13):7367–78. doi: 10.1128/JVI.00554-14. PubMed PMID: 24741095; PubMed Central PMCID: PMCPMC4054434.

29. Sun Z, Jha HC, Robertson ES. Bub1 in Complex with LANA Recruits PCNA To Regulate Kaposi’s Sarcoma-Associated Herpesvirus Latent Replication and DNA Translesion Synthesis. J Virol. 2015;89(20):10206–18. doi: 10.1128/JVI.01524-15. PubMed PMID: 26223641; PubMed Central PMCID: PMCPMC4580184.

30. Cai Q, Xiao B, Si H, Cervini A, Gao J, Lu J, et al. Kaposi’s sarcoma herpesvirus upregulates Aurora A expression to promote p53 phosphorylation and ubiquitylation. PLoS Pathog. 2012;8(3):e1002566. doi: 10.1371/journal.ppat.1002566. PubMed PMID: 22396649; PubMed Central PMCID: PMCPMC3291660.

31. Lu J, Verma SC, Murakami M, Cai Q, Kumar P, Xiao B, et al. Latency-associated nuclear antigen of Kaposi’s sarcoma-associated herpesvirus (KSHV) upregulates survivin expression in KSHV-Associated B-lymphoma cells and contributes to their proliferation. J Virol. 2009;83(14):7129–41. doi: 10.1128/JVI.00397-09. PubMed PMID: 19439469; PubMed Central PMCID: PMCPMC2704763.

32. Si H, Verma SC, Lampson MA, Cai Q, Robertson ES. Kaposi’s sarcoma-associated herpesvirus-encoded LANA can interact with the nuclear mitotic apparatus protein to regulate genome maintenance and segregation. J Virol. 2008;82(13):6734–46. doi: 10.1128/JVI.00342-08. PubMed PMID: 18417561; PubMed Central PMCID: PMCPMC2447046.

33. Brown RR, Huis In ’t Veld PJ, Musacchio A, Straight AF. Structure and function of the centromere and inner kinetochore. Nat Rev Mol Cell Biol. 2026;27(10):729–42. doi: 10.1038/s41580-026-00989-7. PubMed PMID: 42547645.

34. Giunta S, Fukagawa T, O’Neill RJ, Akiyoshi B. 40 years of CENP-A: the foundation of a new era of centromere biology. Chromosoma. 2025;134(1):13. doi: 10.1007/s00412-025-00842-4. PubMed PMID: 41460345.

35. Ariyoshi M, Fukagawa T. An updated view of the kinetochore architecture. Trends Genet. 2023;39(12):941–53. doi: 10.1016/j.tig.2023.09.003. PubMed PMID: 37775394.

36. Hara M, Fukagawa T. Critical Foundation of the Kinetochore: The Constitutive Centromere-Associated Network (CCAN). Prog Mol Subcell Biol. 2017;56:29–57. doi: 10.1007/978-3-319-58592-5_2. PubMed PMID: 28840232.

37. Pesenti ME, Raisch T, Conti D, Walstein K, Hoffmann I, Vogt D, et al. Structure of the human inner kinetochore CCAN complex and its significance for human centromere organization. Mol Cell. 2022;82(11):2113–31 e8. doi: 10.1016/j.molcel.2022.04.027. PubMed PMID: 35525244; PubMed Central PMCID: PMCPMC9235857.

38. Cheeseman IM, Chappie JS, Wilson-Kubalek EM, Desai A. The conserved KMN network constitutes the core microtubule-binding site of the kinetochore. Cell. 2006;127(5):983–97. doi: 10.1016/j.cell.2006.09.039. PubMed PMID: 17129783.

39. Ouzounidis VR, Green M, van Capelle CC, Gebhardt C, Crellin H, Finlayson C, et al. The outer kinetochore components KNL-1 and Ndc80 complex regulate axon and neuronal cell body positioning in the C. elegans nervous system. Mol Biol Cell. 2024;35(6):ar83. doi: 10.1091/mbc.E23-08-0325. PubMed PMID: 38656792; PubMed Central PMCID: PMCPMC11238089.

40. Roberts AJ, Kon T, Knight PJ, Sutoh K, Burgess SA. Functions and mechanics of dynein motor proteins. Nat Rev Mol Cell Biol. 2013;14(11):713–26. doi: 10.1038/nrm3667. PubMed PMID: 24064538; PubMed Central PMCID: PMCPMC3972880.

41. Bamba C, Bobinnec Y, Fukuda M, Nishida E. The GTPase Ran regulates chromosome positioning and nuclear envelope assembly in vivo. Curr Biol. 2002;12(6):503–7. doi: 10.1016/s0960-9822(02)00741-8. PubMed PMID: 11909538.

42. Silverman-Gavrila RV, Wilde A. Ran is required before metaphase for spindle assembly and chromosome alignment and after metaphase for chromosome segregation and spindle midbody organization. Mol Biol Cell. 2006;17(4):2069–80. doi: 10.1091/mbc.e05-10-0991. PubMed PMID: 16481399; PubMed Central PMCID: PMCPMC1415283.

43. de Wolf B, Kops G. Kinetochore Malfunction in Human Pathologies. Adv Exp Med Biol. 2017;1002:69–91. doi: 10.1007/978-3-319-57127-0_4. PubMed PMID: 28600783.

44. Putkey FR, Cramer T, Morphew MK, Silk AD, Johnson RS, McIntosh JR, et al. Unstable kinetochore-microtubule capture and chromosomal instability following deletion of CENP-E. Dev Cell. 2002;3(3):351–65. doi: 10.1016/s1534-5807(02)00255-1. PubMed PMID: 12361599.

45. Huang H, Lampson M, Efimov A, Yen TJ. Chromosome instability in tumor cells due to defects in Aurora B mediated error correction at kinetochores. Cell Cycle. 2018;17(23):2622–36. doi: 10.1080/15384101.2018.1553340. PubMed PMID: 30513041; PubMed Central PMCID: PMCPMC6300107.

46. Rozen R, Sathish N, Li Y, Yuan Y. Virion-wide protein interactions of Kaposi’s sarcoma-associated herpesvirus. J Virol. 2008;82(10):4742–50. doi: 10.1128/JVI.02745-07. PubMed PMID: 18321973; PubMed Central PMCID: PMCPMC2346726.

47. Kaul R, Verma SC, Robertson ES. Protein complexes associated with the Kaposi’s sarcoma-associated herpesvirus-encoded LANA. Virology. 2007;364(2):317–29. doi: 10.1016/j.virol.2007.03.010. PubMed PMID: 17434559; PubMed Central PMCID: PMCPMC4067005.

48. Han C, Niu D, Lan K. Rewriting Viral Fate: Epigenetic and Transcriptional Dynamics in KSHV Infection. Viruses. 2024;16(12). doi: 10.3390/v16121870. PubMed PMID: 39772181; PubMed Central PMCID: PMCPMC11680275.

49. Li S, Tan M, Juillard F, Ponnusamy R, Correia B, Simas JP, et al. The Kaposi Sarcoma Herpesvirus Latency-associated Nuclear Antigen DNA Binding Domain Dorsal Positive Electrostatic Patch Facilitates DNA Replication and Episome Persistence. J Biol Chem. 2015;290(47):28084–96. doi: 10.1074/jbc.M115.674622. PubMed PMID: 26420481; PubMed Central PMCID: PMCPMC4653668.

50. Singh RK, Lang F, Pei Y, Jha HC, Robertson ES. Metabolic reprogramming of Kaposi’s sarcoma associated herpes virus infected B-cells in hypoxia. PLoS Pathog. 2018;14(5):e1007062. doi: 10.1371/journal.ppat.1007062. PubMed PMID: 29746587; PubMed Central PMCID: PMCPMC5963815.

51. Purushothaman P, Dabral P, Gupta N, Sarkar R, Verma SC. KSHV Genome Replication and Maintenance. Front Microbiol. 2016;7:54. doi: 10.3389/fmicb.2016.00054. PubMed PMID: 26870016; PubMed Central PMCID: PMCPMC4740845.

52. Losay VA, Damania B. Unraveling the Kaposi Sarcoma-Associated Herpesvirus (KSHV) Lifecycle: An Overview of Latency, Lytic Replication, and KSHV-Associated Diseases. Viruses. 2025;17(2). doi: 10.3390/v17020177. PubMed PMID: 40006930; PubMed Central PMCID: PMCPMC11860327.

53. Zhang F, Liang D, Lin X, Zou Z, Sun R, Wang X, et al. NDRG1 facilitates the replication and persistence of Kaposi’s sarcoma-associated herpesvirus by interacting with the DNA polymerase clamp PCNA. PLoS Pathog. 2019;15(2):e1007628. doi: 10.1371/journal.ppat.1007628. PubMed PMID: 30811506; PubMed Central PMCID: PMCPMC6411202.

54. Sorel O, Dewals BG. The Critical Role of Genome Maintenance Proteins in Immune Evasion During Gammaherpesvirus Latency. Front Microbiol. 2018;9:3315. doi: 10.3389/fmicb.2018.03315. PubMed PMID: 30687291; PubMed Central PMCID: PMCPMC6333680.

55. Cohen JI. Herpesvirus latency. J Clin Invest. 2020;130(7):3361-9. doi: 10.1172/JCI136225. PubMed PMID: 32364538; PubMed Central PMCID: PMCPMC7324166 exists.

56. Speck SH, Ganem D. Viral latency and its regulation: lessons from the gamma-herpesviruses. Cell Host Microbe. 2010;8(1):100–15. doi: 10.1016/j.chom.2010.06.014. PubMed PMID: 20638646; PubMed Central PMCID: PMCPMC2914632.

57. Suntronpong A, Kugou K, Masumoto H, Srikulnath K, Ohshima K, Hirai H, et al. CENP-B box, a nucleotide motif involved in centromere formation, occurs in a New World monkey. Biol Lett. 2016;12(3):20150817. doi: 10.1098/rsbl.2015.0817. PubMed PMID: 27029836; PubMed Central PMCID: PMCPMC4843215.

58. Chardon F, Japaridze A, Witt H, Velikovsky L, Chakraborty C, Wilhelm T, et al. CENP-B-mediated DNA loops regulate activity and stability of human centromeres. Mol Cell. 2022;82(9):1751–67 e8. doi: 10.1016/j.molcel.2022.02.032. PubMed PMID: 35320753.

59. Nagpal H, Ali-Ahmad A, Hirano Y, Cai W, Halic M, Fukagawa T, et al. CENP-A and CENP-B collaborate to create an open centromeric chromatin state. Nat Commun. 2023;14(1):8227. doi: 10.1038/s41467-023-43739-5. PubMed PMID: 38086807; PubMed Central PMCID: PMCPMC10716449.

60. Kumari A, Kumar C, Wasnik N, Mylavarapu SVS. Dynein light intermediate chains as pivotal determinants of dynein multifunctionality. J Cell Sci. 2021;134(10). doi: 10.1242/jcs.254870. PubMed PMID: 34014309.

61. Mische S, He Y, Ma L, Li M, Serr M, Hays TS. Dynein light intermediate chain: an essential subunit that contributes to spindle checkpoint inactivation. Mol Biol Cell. 2008;19(11):4918–29. doi: 10.1091/mbc.e08-05-0483. PubMed PMID: 18799620; PubMed Central PMCID: PMCPMC2575169.

62. Singh RK, Bose D, Robertson ES. Epigenetic Reprogramming of Kaposi’s Sarcoma-Associated Herpesvirus during Hypoxic Reactivation. Cancers (Basel). 2022;14(21). doi: 10.3390/cancers14215396. PubMed PMID: 36358814; PubMed Central PMCID: PMCPMC9654037.

63. Singh RK, Bose D, Robertson ES. HIF1alpha-Regulated Expression of the Fatty Acid Binding Protein Family Is Important for Hypoxic Reactivation of Kaposi’s Sarcoma-Associated Herpesvirus. J Virol. 2021;95(12). doi: 10.1128/JVI.02063-20. PubMed PMID: 33789996; PubMed Central PMCID: PMCPMC8316119.

64. Kumar Singh R, Pei Y, Bose D, Lamplugh ZL, Sun K, Yuan Y, et al. KSHV-encoded vCyclin can modulate HIF1alpha levels to promote DNA replication in hypoxia. Elife. 2021;10. doi: 10.7554/eLife.57436. PubMed PMID: 34279223; PubMed Central PMCID: PMCPMC8315796.

65. Verma SC, Lu J, Cai Q, Kosiyatrakul S, McDowell ME, Schildkraut CL, et al. Single molecule analysis of replicated DNA reveals the usage of multiple KSHV genome regions for latent replication. PLoS Pathog. 2011;7(11):e1002365. doi: 10.1371/journal.ppat.1002365. PubMed PMID: 22072974; PubMed Central PMCID: PMCPMC3207954.

